# Noise propagation shapes condition-dependent gene expression noise in *Escherichia coli*

**DOI:** 10.1101/795369

**Authors:** Arantxa Urchueguía, Luca Galbusera, Gwendoline Bellement, Thomas Julou, Erik van Nimwegen

## Abstract

Although it is well appreciated that gene expression is inherently noisy and that transcriptional noise is encoded in a promoter’s sequence, little is known about the variation in transcriptional noise across growth conditions. Using flow cytometry we here quantify transcriptional noise in *E. coli* genome-wide across 8 growth conditions, and find that noise and gene regulation are intimately coupled. Apart from a growth-rate dependent lower bound on noise, we find that individual promoters show highly condition-dependent noise and that condition-dependent expression noise is shaped by noise propagation from regulators to their targets. A simple model of noise propagation identifies TFs that most contribute to both condition-specific and condition-independent noise propagation. The overall correlation structure of sequence and expression properties of *E. coli* genes uncovers that genes are organized along two principal axes, with the first axis sorting genes by their mean expression and evolutionary rate of their coding regions, and the second axis sorting genes by their expression noise, the number of regulatory inputs in their promoter, and their expression plasticity.

## Introduction

It is by now well established that isogenic cells growing in a homogeneous environment show cell-to-cell fluctuations in gene expression, e.g. Elowitz *et al*, 2002; Rao *et al*, 2002; Blake *et al*, 2003; Raser and O’Shea, 2005. This gene expression noise is not surprising from a biophysical perspective given the inherent thermodynamic fluctuations in the molecular events underlying gene expression and the small numbers of molecules involved. Much progress has been made in understanding the sources and mechanisms underlying this stochastic heterogeneity in gene expression, including fluctuations in transcription and translation, chromatin state, or cellular growth, e.g. Golding *et al*, 2005; Blake *et al*, 2006; Tirosh and Barkai, 2008; Raj and van Oudenaarden, 2008; Sanchez *et al*, 2013; Carey *et al*, 2013; Shahrezaei and Marguerat, 2015; Keren *et al*, 2015.

From a functional perspective, stochastic heterogeneity in gene expression, or more generally in phenotypic state, has generally been seen as a mechanism that is complementary to gene regulation. That is, whereas gene regulatory mechanisms allow cells to sense and respond to changes in their environment in a targeted manner, stochastic heterogeneity provides an alternative ‘bet hedging’ strategy enabling cells to deal with varying environments, e.g. Bull, 1987; Haccou and Iwasa, 1995; Kussell and Leibler, 2005; Tanase-Nicola and ten Wolde, 2008; Ackermann *et al*, 2008.

However, in recent years evidence has been accumulating that gene regulation and gene expression noise may be inherently and intimately linked. For example, several studies have shown that transcriptional noise varies significantly across genes and is to a substantial extent encoded in the promoter sequence of a gene Taniguchi *et al*, 2010; Silander *et al*, 2012; Carey *et al*, 2013; Jones *et al*, 2014; Wolf *et al*, 2015. Indeed, whereas genes that are transcribed at a constant rate will exhibit Poissonian fluctuations in mRNA levels, most genes exhibit significantly higher levels of transcriptional noise. This increased transcriptional noise is generally understood to result, at least partially, from the fact that binding and unbinding of transcription factors (TFs) causes the promoter to stochastically switch between different states that are associated with different transcription initiation rates. In this way, fluctuations in both the expression levels of TFs and their binding to promoter regions are propagated to the their target genes and this *noise propagation* has long been recognized as an unavoidable side effect of regulation Thattai and van Oudenaarden, 2001; Pedraza, 2005; Lestas *et al*, 2010; Lehner and Kaneko, 2011; Bruggeman and Teusink, 2018.

In a recent study we showed that, in *E. coli*, unregulated promoters have low expression noise by default, and that the more regulatory inputs a gene has, the more noisy its gene expression tends to be Wolf *et al*, 2015. A similar general association between highly regulated genes and high expression noise has also been observed in eukaryotes Blake *et al*, 2003; Newman *et al*, 2006, and it has also been observed that co-regulated genes show correlated gene expression fluctuations Junker and van Oudenaarden, 2012. All these observations suggest that noise propagation may be a key determinant of gene expression noise.

If noise propagation is indeed an important determinant of gene expression noise, then expression noise should not be an intrinsic property of a gene, but should be *condition-dependent*. That is, fluctuations in a promoter’s transcription rate depend on the stochastic binding and unbinding of TFs and these in turn depend on average expression levels and fluctuations in expression levels across cells Becskei *et al*, 2005; Carey *et al*, 2013; Sharon *et al*, 2014; Jones *et al*, 2014. Thus, as TFs change their expression levels across conditions, the noise properties of their target promoters should vary as well. However, so far there has been no systematic investigation into how the noise properties of genes in *E. coli* vary across conditions.

We here systematically quantify how genome-wide gene expression noise in *E. coli* varies across conditions by using flow cytometry in combination with a library of fluorescent transcriptional reporters Zaslaver *et al*, 2006 in 8 different growth conditions, including different nutrients, stresses, and in stationary phase. We investigate how global noise properties vary across conditions and quantify noise propagation by modeling the condition-dependent transcriptional noise of each gene in terms of annotated regulatory sites in their promoters. Using this modeling we infer which TFs are contributing most to expression noise in each condition, and identify several TFs that consistently contribute to noise propagation in all growth conditions. Our analysis shows that gene expression noise and gene regulation are intimately coupled. In particular, the number of regulatory inputs of a gene, its expression plasticity, its gene expression noise, and also the plasticity in its gene expression noise, are all highly positively correlated.

## Results

### Signatures of noise propagation

Understanding how noise propagation from TFs shapes patterns of gene expression noise is experimentally challenging as it is difficult to manipulate the activity of TFs in a predictable way. However, the activities of TFs are generally expected to change between conditions. Therefore, if noise propagation is a key determinant of gene expression noise, we expect noise levels across different conditions to exhibit certain qualitative features, as illustrated in Fig. 1.

**Figure 1:**
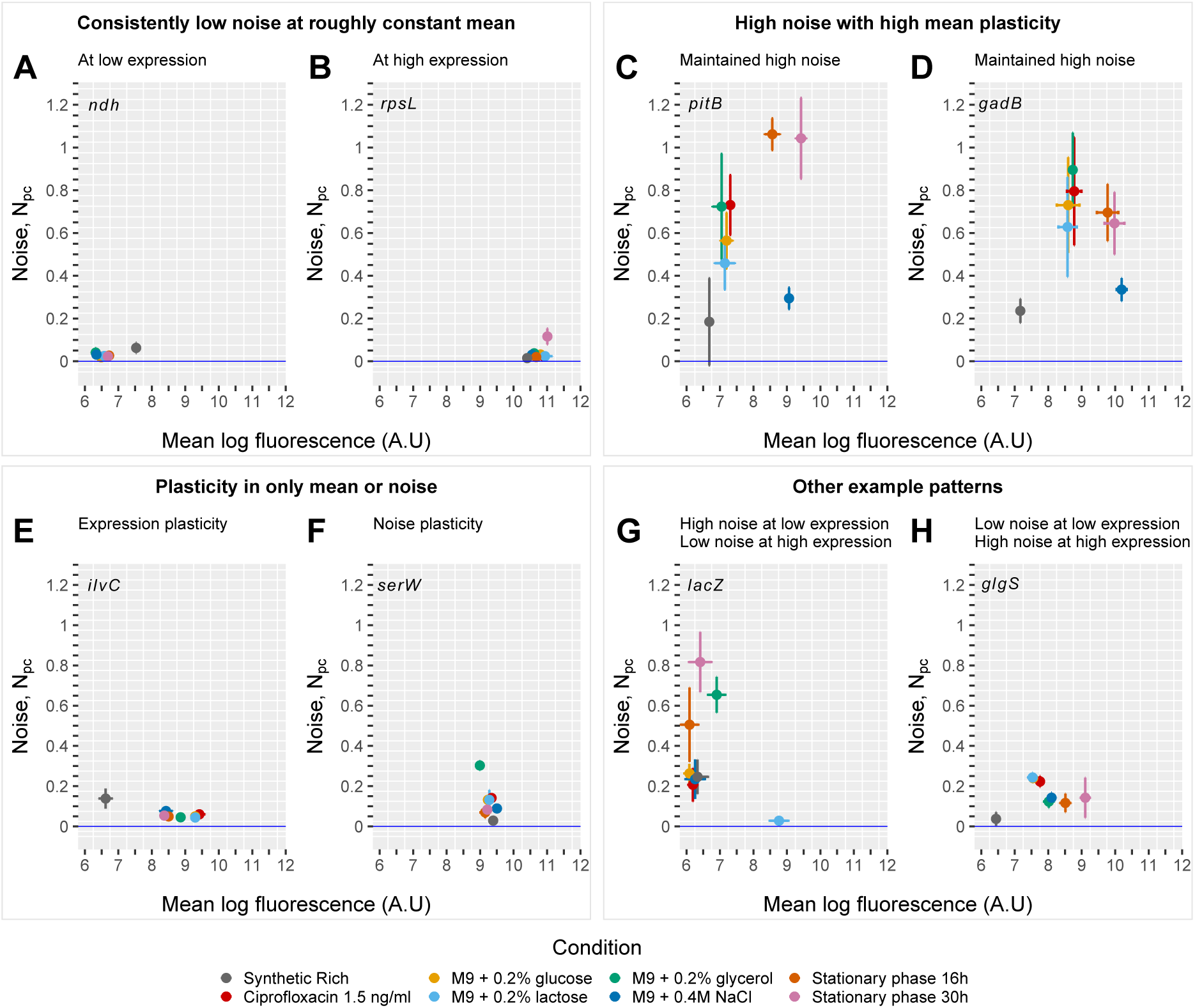
Condition-dependent noise propagation. One mechanism by which noise propagation can arise is from heterogeneous expression of transcription factors. **A:** Two independent transcription factors (TF) represented each by an orange and a blue dot show differential single-cell expression distributions across two conditions. TF1 (blue dots) is homogeneously expressed in condition 1, but not in condition 2, whereas TF2 (orange dots) shows the opposite behaviour. **B:** As noise propagation is a condition-dependent mechanism, in the condition where a given transcription displays a heterogeneous expression, its targets will also show increased variability. As shown in the illustration, since TF2 is noisier than TF1 in condition 1, its targets show also higher noise than those of TF1, whereas in condition 2 the opposite occurs. **C:** Given the condition-dependent nature of noise propagation, regulated promoters will show larger plasticity in noise across conditions than constitutive ones, as the latter are not affected by condition-dependent noise propagation. Here we define noise plasticity as the variance in noise levels of a promoter across conditions.

We consider a simple case scenario where two individual transcription factors show variable activities, i.e. expression levels and DNA binding, both across conditions and across cells within each condition. In condition 1, we assume TF 1 to be higher expressed on average and less variable in expression across cells than TF 2, whereas in condition 2 the situation is reversed, i.e. TF 2 has higher mean and less variability across cells (Fig. 1A). As a consequence of noise propagation, the cumulative distribution of noise levels for the targets of TF 2 will be shifted to higher values compared to the targets of TF 1 in condition 1, whereas in condition 2 the situation will be reversed (Fig. 1B). In other words, targets of TFs that change their expression distribution across conditions are expected to change their relative noise levels across conditions. In addition, by comparing the relative noise levels of targets of different TFs, it should be possible to infer which TFs are contributing most to noise propagation in a giving condition, and below we develop a model approach to do this. Finally, because constitutively expressed promoters are not affected by noise propagation, they are expected to change their noise levels less across conditions than regulated genes that are subject to noise propagation. That is, regulated genes are expected to have higher *noise plasticity* (Fig. 1C).

### Expression noise levels vary substantially across conditions and on average decrease with growth-rate

We used flow cytometry together with a library of fluorescent transcriptional reporters to measure gene expression distributions of *E. coli* promoters genome-wide across a set of 8 different growth conditions (Fig. 2A). The library (Zaslaver *et al*, 2006) consists of most of *E. coli*’s intergenic regions inserted upstream of a strong ribosomal binding site and a fast-folding GFP in a low copy plasmid, and has already been used in several studies to study gene expression noise in *E.coli* Freed *et al*, 2008; Silander *et al*, 2012; Wolf *et al*, 2015. As we have shown previously Wolf *et al*, 2015, GFP levels of these reporters reflect transcriptional activity, since translation and mRNA decay rates vary little across reporters because their mRNAs are almost identical.

**Figure 2:**
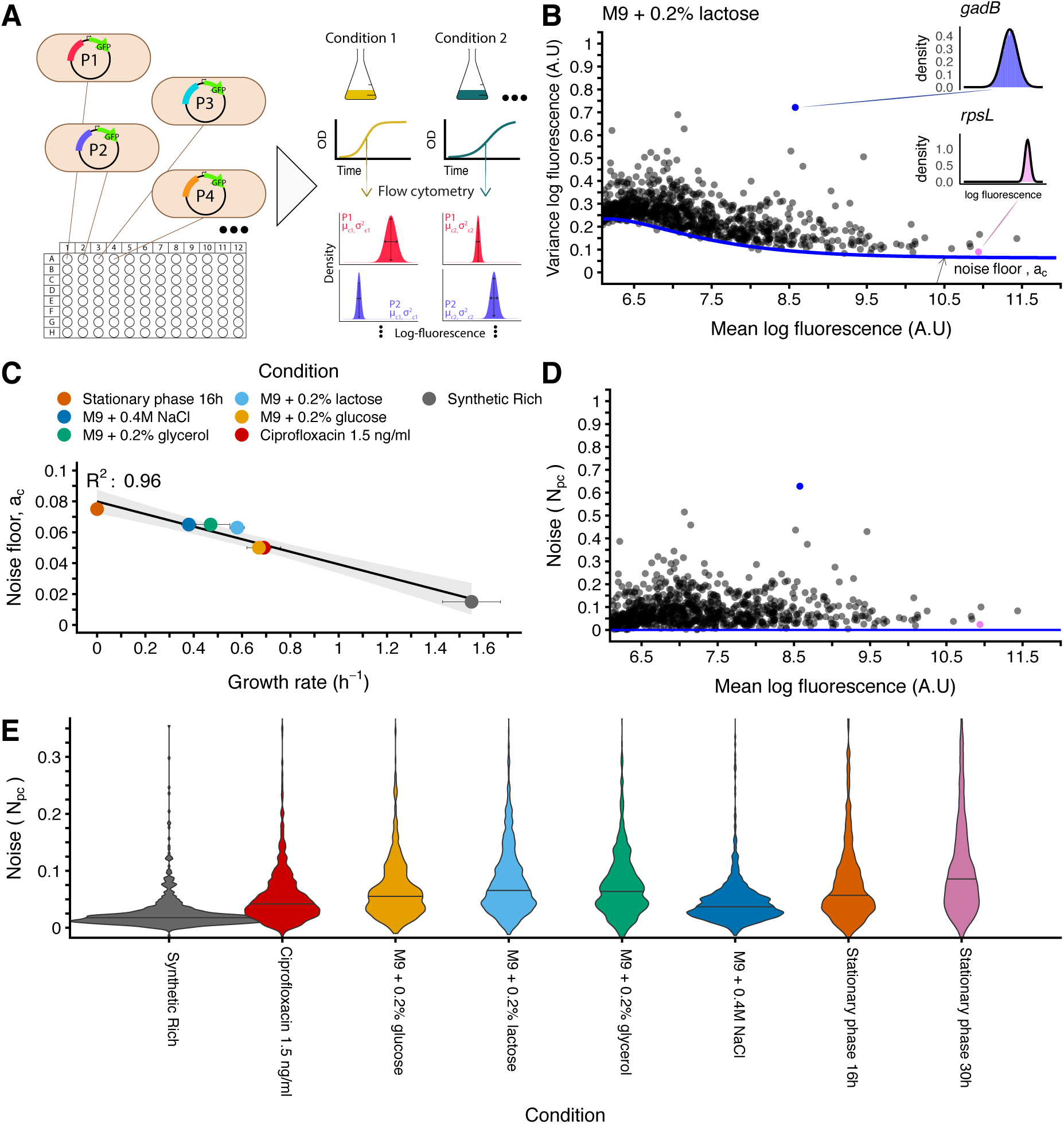
Expression noise of native *E. coli* promoters under different growth conditions. **A:** For each growth condition and *E. coli* promoter, we used flow cytometry to measure the distribution of GFP levels across single cells of the corresponding fluorescent reporter. The 8 growth conditions comprised synthetic rich media, minimal media with different carbon sources, an osmotic and DNA damage stress, and two time points in stationary phase. **B:** Mean (x-axis) and variance (y-axis) of log GFP levels for all promoters with expression above a background level for growth in M9 0.2% lactose (see Suppl. Fig. S7 for results in all conditions). The blue line shows the fitted minimal variance as a function of mean expression and the corresponding noise floor *a*_*c*_ is indicated with an arrow. The insets show distributions of log-GFP levels for two example promoters. **C:** The noise floor *a*_*c*_ as a function of the growth rate in the respective condition (*stationary phase at 30h not shown*). The line indicates a linear fit (with 0.95 confidence interval in grey) and the Pearson squared correlation coefficient *R*^2^ is also indicated. **D:** To compare noise of promoters with different means, we defined the noise level of a promoter as the difference between its variance and the fitted minimal variance at its mean expression. Shown are noise levels versus mean for promoters in M9 0.2% lactose. **E:** Noise level distributions of the full library in each of the measured conditions. The horizontal lines indicate the medians. The vertical scale is clipped at 0.35 for better visibility (Suppl. Fig. S8 has the full distributions).

The growth conditions (Suppl. Table S1) were chosen to span a wide range of growth rates (Suppl. Fig. S1), cell physiologies (Suppl. Fig. S2), and regulatory states. They consist of M9 minimal media with three different carbon sources (0.2% glucose, 0.2% glycerol and 0.2% lactose), two stresses (sub-MIC antibiotic: Ciprofloxacin 1.5 ng/ml + 0.2% glucose and osmotic: 0.4M NaCl + 0.2% glucose), two time points in stationary phase (after 16h and 30h of growth in 0.2% glucose, respectively), and a MOPS synthetic rich media. We used microscopy to image cells from each growth condition and found that, consistent with the known relationship between growth-rate and cell physiology Schaechter *et al*, 1958, cell size generally increased with growth-rate (Suppl. Fig. S3).

For each condition and each promoter, we used high-throughput flow cytometry to measure GFP levels for thousands of single-cells. Apart from the two stationary phase conditions, all measurements were taken during mid-exponential phase. In total we gathered 50^*1*^000 single-cell measurements for each of the 1810 promoters in the library across 8 conditions, including some conditions in replicate. As observed previously Wolf *et al*, 2015, the fluorescence distributions can be well fitted with log-normal distributions and we thus characterized each fluorescence distribution by the mean and variance of log-fluorescence. To estimate mean and variance we used a method that uses forward and side scatter to identify viable cells and fits the log-fluorescence distribution by a mixture of a Gaussian and uniform distribution to remove possible outliers (e.g. contaminants, non-growing cells) Galbusera *et al*, 2019.

Replicate measurements performed on different days were highly reproducible, with Pearson correlations *R*^2^ *>* 0.99 for the mean between replicates in all conditions, and correlations for the variance ranging from *R*^2^ = 0.85 to *R*^2^ = 0.95 (Suppl. Fig. S4). In order to determine whether the variability derived mainly from biological variation from day-to-day or from measurement noise, we performed a time-course experiment where we repeatedly measured the same culture at different time points during exponential growth and found that both the mean and variance measurements were extremely reproducible (Suppl. Figs. S5 and S6). This indicates that the differences we observed among replicates from different days mainly reflect biological variability and not technical measurement errors. This also implies that genes exhibit more biological variation in their noise levels across days than in their mean expression.

As an example, Fig. 2B shows the variance as a function of mean for each promoter measured in M9 minimal media + 0.2% lactose (see Suppl. Fig. S7 for all conditions). Note that the variance in log-fluorescence is equal to the square of the coefficient of variation (*CV* ^2^) whenever fluctuations are small relative to the mean Wolf *et al*, 2015. This approximation applies in our data, as the majority of promoters (~75% across all conditions) have a variance smaller than 0.3 (Suppl. Fig. S7).

As has been observed in previous studies Bar-Even *et al*, 2006; Newman *et al*, 2006; Taniguchi *et al*, 2010; Wolf *et al*, 2015, there is a clear lower bound on noise as a function of the mean expression level of the promoter (Fig. 2B). As explained in the Material and Methods section, we have previously shown Wolf *et al*, 2015 that the functional form of the minimal variance as a function of mean expression, equation (3), can be derived assuming that GFP variance is the sum of two terms: one ‘multiplicative’ contribution with variance proportional to the square of the mean expression, and one ‘Poissonian’ contribution with variance proportional to mean expression. The former term, which we will refer to as the ‘noise floor’ *a*_*c*_, corresponds to the minimal variance for highly expressed promoters and likely results from global fluctuations in transcription, translation, mRNA decay, and growth Elowitz *et al*, 2002; Taniguchi *et al*, 2010. This term is often referred to as an ‘extrinsic noise’ contribution. The Poissonian term, whose magnitude we denote by *b*_*c*_ and is often referred to as the ‘intrinsic noise’ term, could in principle derive from intrinsic expression noise whose magnitude scales proportional to mean expression Taniguchi *et al*, 2010; Sánchez and Kondev, 2008. However, by comparing microscopy and flow cytometry measurements we have recently shown that, at these expression levels, the component *b*_*c*_ derives almost entirely from the measurement noise of the flow cytometer Galbusera *et al*, 2019. As shown in Suppl. Fig. S7, the same functional form describes the minimal variance in all conditions and we estimated the noise floor *a*_*c*_ at high expression in each condition.

Remarkably, we observed that the noise floor *a*_*c*_ is an almost perfectly decreasing linear function of growth-rate (*R*^2^ = 0.96, Fig. 2C). Thus, the slower cells grow, the higher the minimal cell-to-cell variability in gene expression. A similar anti-correlation between noise and growth-rate has been previously observed in eukaryotes, but was proposed to derive from heterogeneity in cell cycle stage Keren *et al*, 2015. However, our results show that this general anti-correlation between noise and growth-rate also occurs in prokaryotes that do not have analogous cell cycle stages.

In order to have a noise measure for each gene that does not systematically depend on its mean expression, we defined the noise level *N*_*pc*_ of promoter *p* in condition *c* as the difference between its variance in log-fluorescence and the minimal variance at its mean expression level (Materials and Methods equation (4) and Fig. 2D). Figure 2E shows the distribution of noise levels *N*_*pc*_ in each of the conditions, sorted from high to low growth-rate. We see that not only the noise floor, but also the distribution of noise on levels on top of this noise floor varies substantially across conditions. Moreover, like the noise floor, both the median of the noise levels *N*_*pc*_ as well as the variability in noise levels increase as the growth-rate decreases, e.g. the noise levels are lowest in synthetic rich conditions (*p* = 3 × 10^*−*30^, Wilcoxon rank-sum test) and highest at 30h of stationary phase (*p* = 5 × 10^*−*68^, Wilcoxon rank sum test). That is, not only do minimal noise levels increase as growth-rate decreases, the variability in noise levels across genes increases as well. The only exception to this general trend is the osmotic stress condition M9 + 0.4M NaCl, which has relatively low variability in noise levels *N*_*pc*_ compared to other conditions with similar growth-rate (Fig. 2E), even though its noise floor is *not* deviating from the general dependence on growth-rate. These results show that growth-rate, and more generally the physiological state of the cell has a major influence on the absolute noise levels. However, in this work we will focus on how the *relative* noise levels of different promoters vary across conditions.

### Individual promoters show highly diverse changes in noise across conditions

If changes in noise levels across conditions were mostly driven by the overall physiology of the cells, then we would expect different genes to exhibit coherent changes in noise across conditions. For example, noise levels might rescale across conditions as a function of the mean expression of the gene in the condition. In contrast, we observe that different promoters show highly diverse changes in their noise levels across conditions (Fig. 3). Some promoters show consistently low noise at either low or high mean expression (Fig. 3A and B), some promoters show consistently high noise that also strongly varies across conditions in a manner not correlated with mean expression (Fig. 3C and D), some promoters show only plasticity in mean (Fig. 3E) or only plasticity in noise (Fig. 3F), but many other patterns of behavior were observed. For example, there are also promoters that show only low noise when the promoter has high mean (Fig. 3G), or only low noise when the promoter has low mean (Fig. 3H).

**Figure 3:**
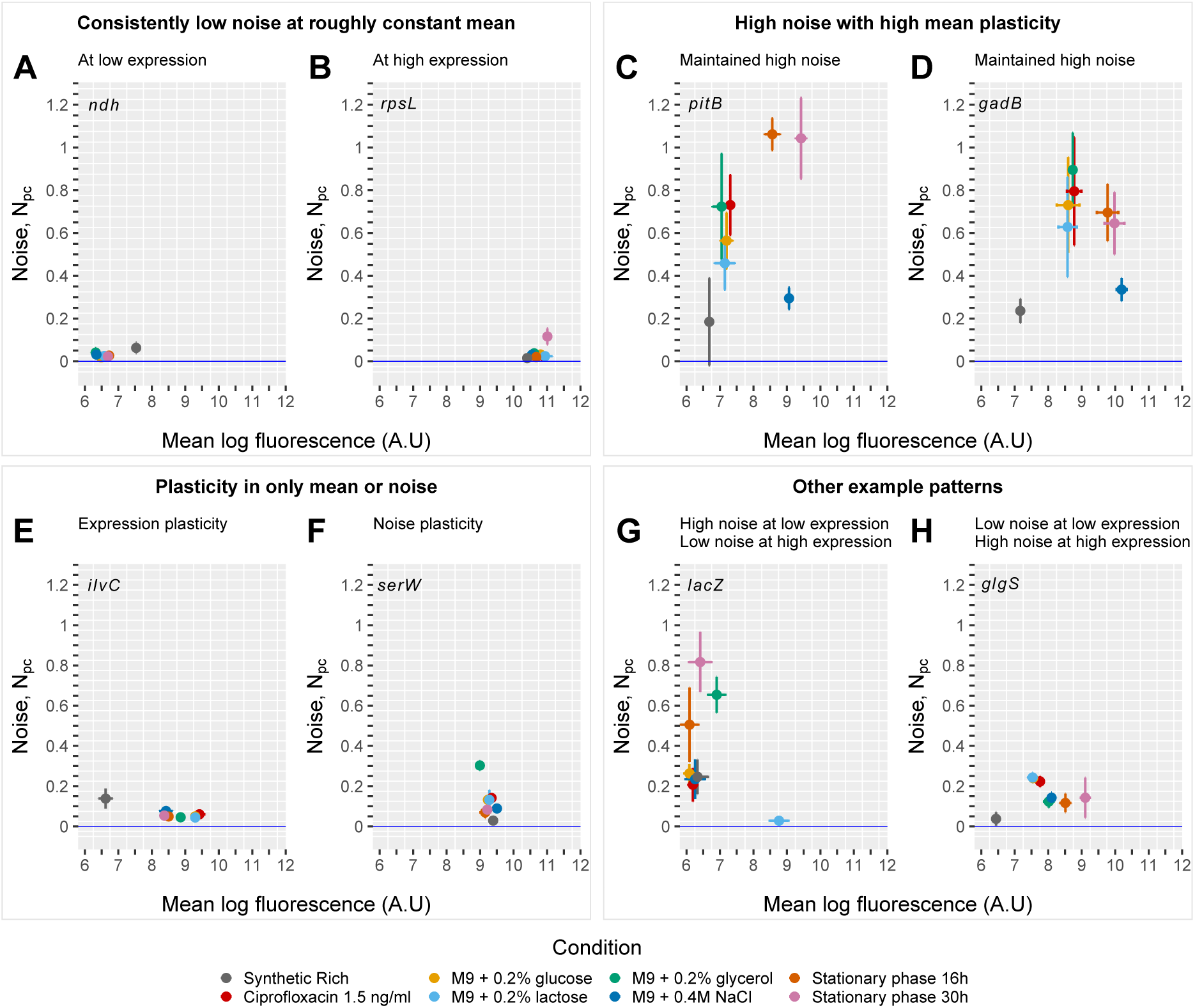
Individual promoters show diverse patterns of changes in noise levels across conditions. Each panel shows the noise level as a function of mean across conditions (different colors) for an individual promoter (in each panel the name of the immediately downstream gene is indicated). Error bars denote standard-errors of the estimates. Each of the 4 pairs of panels indicate different types of behavior in mean and noise across conditions, as described at the top of each pair of panels.

The growth media was not a predictor of how individual genes were going to change their mean and noise. For example, while overall the whole library is shifted towards lower noise in synthetic rich media, individual genes can show higher noise in this condition compared to other conditions (e.g. Fig. 3A, E, and G). We highlighted this particular condition as an example, but the same observation applies to others. These observations indicate that global changes in the cell physiology or in the expression level only cannot explain how the noise of a promoter varies across conditions. This implies that there is a promoter specific source of noise shaping gene expression variability across the environments.

### Noise propagation explains the condition-dependent noise levels of genes

That noise propagation from regulators to their targets may play an important role in setting the relative noise levels across genes was suggested by previous analysis Wolf *et al*, 2015, where we found there to be a substantial correlation between the noise level of a gene in M9 + glucose and the number of regulatory inputs it has, i.e. the number of different TFs that are known to regulate it according to RegulonDB Santos-Zavaleta *et al*, 2019. Here we find that this positive association between the noise level of a gene and its number of regulatory inputs is observed in all 8 conditions that we tested (Fig. 4A, Suppl. Fig. S9, Material and Methods). In addition, since the noise levels of TFs likely vary across conditions (Fig. 1C), we expect genes with many regulatory inputs to also show larger noise plasticity, i.e. higher variance in their noise levels across conditions. As Fig. 4B shows, we indeed observe that genes with more regulatory inputs show larger noise plasticity compared to genes with few regulatory inputs or unregulated genes (*p* < 3.7 × 10^*−*10^, two-sided Welch’s t-test).

**Figure 4:**
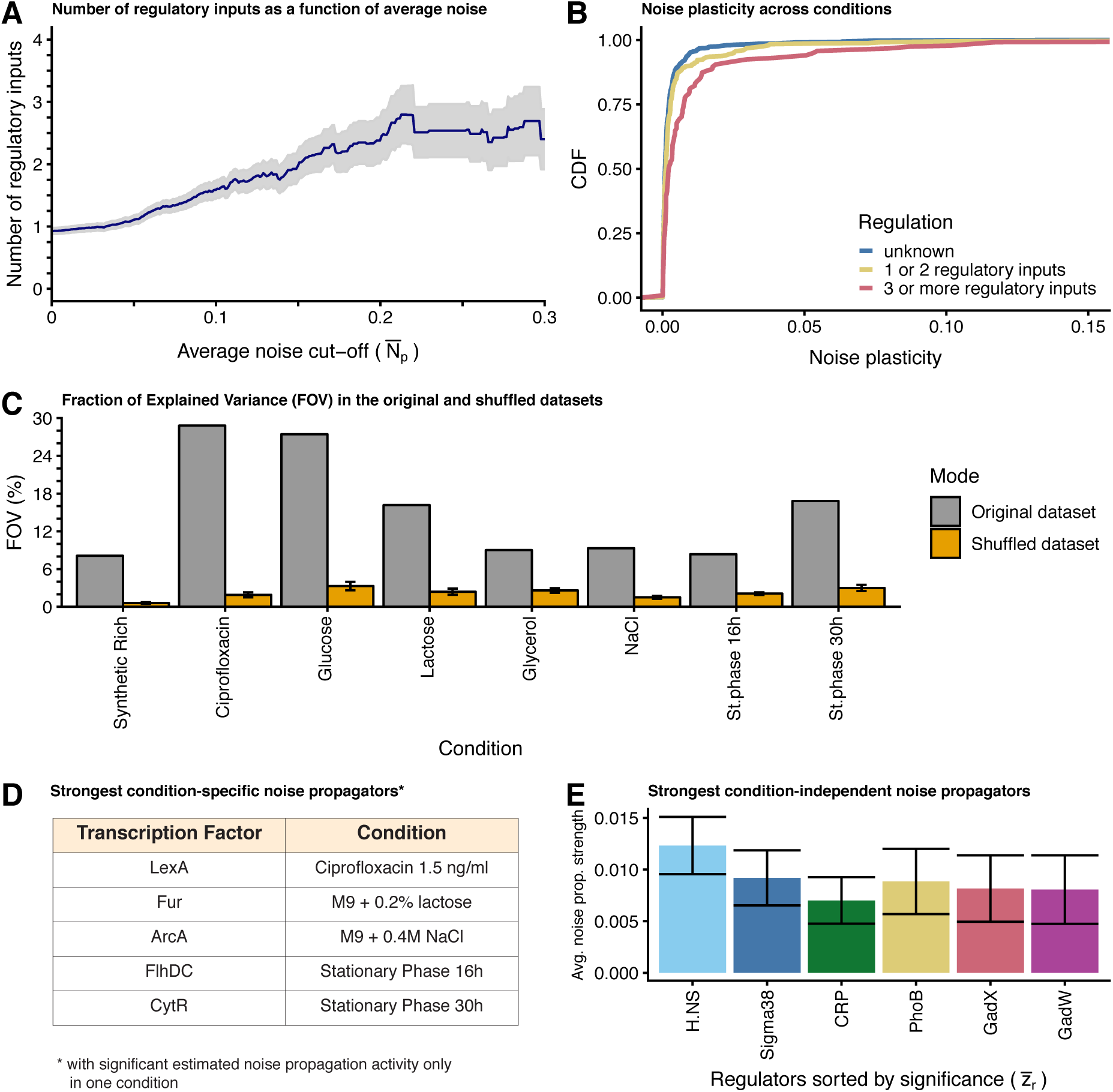
Condition-dependent noise propagation features. **A:** There is a positive association between number of regulatory inputs and noise level. We sorted promoters by their average noise 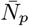 across the 8 conditions and calculated the mean (y-axis) and standard error (grey area) of the average number of known unique regulatory inputs of all promoters with noise above 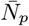, as a function of 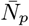 (x-axis). **B:** Promoters with many regulatory inputs show larger noise plasticity than unregulated ones. Shown is the cumulative distribution of the variance in noise of each promoter across the 8 conditions for promoters without known regulatory inputs (blue), 1 or 2 known regulators (yellow), and 3 or more known regulators (red). **C:** Fraction of Explained Variance (FOV, %) by the adapted Motif Activity Response Analysis model (y-axis) in each of the 8 conditions (x-axis) after running it in two modes: on the original dataset (grey bars) and on a randomizeddataset (yellow bars). Randomized data were generated by shuffling the association between regulatory inputs and expression noise multiple times and shown is the average FOV value obtained +/− standard error. **D:** Table of transcription factors predicted by the model as significant condition-specific noise propagators (with *A*_*rc*_ > *δA*_*rc*_). **E:** Average noise propagation strengths (*Ā*_*r*_, y-axis) and their error bars (*δĀ*_*r*_, vertical lines) of the strongest 6 noise-propagators (with *Ā*_*r*_ > *δĀ*_*r*_), sorted by significance (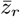, x-axis), that consistently contribute to explain noise levels in all 8 conditions.

If noise propagation is a key determinant of the condition-dependent changes in the noise levels of promoters, then it should be possible to explain some of these changes in terms of the regulatory sites occurring their promoter sequences. We have previously developed a model, called Motif Activity Response Analysis Suzuki *et al*, 2009; Balwierz *et al*, 2014 that models gene expression patterns in terms of computationally predicted regulatory sites in promoters genome-wide and ‘activities’ of regulators. Here we adapted this approach to model the condition-dependent noise levels of promoters in terms of known regulatory inputs and ‘noise propagating activities’ of regulators. In particular, we model the noise *N*_*pc*_ of each promoter *p* in each condition *c* as a linear function of its known regulatory inputs *S*_*pr*_, and the unknown noise propagating activities *A*_*rc*_ of each regulator *r* in each condition *c*:

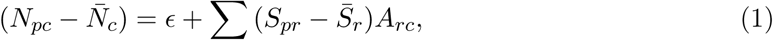

where 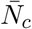 is the average noise level of all promoters in condition *c*, and *ϵ* is a noise term that is assumed Gaussian distributed with mean 0 and unknown variance. We used the RegulonDB database Santos-Zavaleta *et al*, 2019 to set a binary matrix of known regulatory inputs, i.e. *S*_*pr*_ is 1 when promoter *p* is known to be regulated by TF *r* and 0 otherwise. In addition 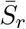 is the average of *S*_*pr*_ across all promoters, i.e. the fraction of promoters targeted by regulator *r*.

For each condition *c* we infer the noise propagating activities *A*_*rc*_ by fitting the model (1) using a Gaussian prior on the activities *A*_*rc*_ to avoid overfitting, which allows us to calculate a full posterior probability distribution over the activities *A*_*rc*_ Balwierz *et al*, 2014. It should be noted that this extremely simple linear model of course only provides a caricature of the complex interactions between TFs and promoters, i.e. we ignore the number, positioning, and affinities of the binding sites, the potential interactions between binding sites for different TFs, the non-linear dependence on TF concentrations, and so on. We thus do not expect that this simple model will accurately predict the expression noise of each promoter across the conditions. Rather, the main aim is to test whether noise propagation can explain a significant fraction of the variation in noise levels across promoters, and to identify which TFs are most responsible for noise propagation in each condition.

As shown in Fig. 4C (grey bars), a substantial fraction of the variance in noise levels in each condition, i.e. between 9% and 29%, can be explained by the simple model of equation (1). To confirm the significance of these fits we fitted the same model to data in which rows of the noise matrix *N*_*pc*_ were randomly shuffled, i.e. the association between regulatory inputs and noise levels were randomized, and observed that the fraction of explained variance on the randomized data was always much lower (Fig. 4C, yellow bars).

Apart from estimating the noise propagating activities *A*_*rc*_ of each regulator *r* in each condition *c*, the model (1) calculates an error bar *δA_rc_* for each of these activities and Suppl. Fig. S10 shows, for each condition, all TFs for which the noise propagating activity was larger than its error-bar, i.e. *A*_*rc*_ > *δA*_*rc*_. We first focused on TFs that contributed to noise-propagation in a highly condition-specific manner. As shown in Fig. 4D, there were 5 TFs that had significant noise propagating activity in only 1 condition. For example, the TF LexA contributed to noise propagation only in the sub-MIC ciprofloxacin condition. LexA is a repressor of the SOS response genes which responds to DNA damage by auto-cleaving on recA polymers that form on DNA double-strand breaks Giese *et al*, 2008. Indeed, it is known ciprofloxacin can induce DNA damage and induce the SOS response Phillips *et al*, 1987. In particular, since we employed ciprofloxacin at a concentration well below the minimal inhibitory concentration, DNA damage likely only occurred in a subset of the cells, leading to heterogeneity in lexA activity across the cells. A second example of a condition-specific noise propagating TF is ArcA, whose activity was only significant in M9 + 0.4M NaCl. ArcA is a general regulator that controls the aerobic/anaerobic expression of respiratory proteins and diverges metabolism into fermentation Salmon *et al*, 2005. It is known that under salt stress major adaptations in metabolism occur and fermentation products increase Arense *et al*, 2010, which is consistent with heterogeneous activity of ArcA in these conditions. Third, we found that FlhDC, the master regulator of flagellar biosynthesis Guttenplan and Kearns, 2013, was contributing to noise propagation only in early stationary phase, i.e after 16h of growth on 0.2% glucose. It is known that flagellar synthesis has a peak in expression during late exponential phase and decreases shortly after entry in stationary phase Amsler *et al*, 1993. Since the 16h condition is a transition between late exponential growth and entry into stationary phase, it seems plausible that some cells had entered growth arrest and were no longer expressing components of the flagellar machinery, while others had not yet transitioned. Fourth, the TF CytR was found to contribute to noise propagation only in late stationary phase. CytR regulates genes involved in nucleoside uptake and utilization Sernova and Gelfand, 2012 and it was recently found that mutations in CytR have a fitness advantage during long term stationary phase Kram *et al*, 2017, which was hypothesized to result from an increased ability to import and use nucleosides that occur in the stationary phase environment due to cell death. Heterogeneity in CytR activation late in stationary phase is consistent with this functional role. Finally, the TF Fur, which regulates genes involved in iron homeostasis Vassinova and Kozyrev, 2000, had significant noise propagating activity only in the M9 + 0.2% lactose condition. In contrast to the other four cases, we do not have a obvious biological interpretation for why Fur activity might be especially heterogeneous during growth on lactose.

In addition to these condition-specific noise propagators, it is noteworthy that many of the most significant noise propagators were found in multiple conditions (Suppl. Fig. S10). To identify regulators that were consistently contributing to noise propagation in all conditions we calculated, for each regulator *r*, its average noise propagating activity *Ā*_*r*_ averaged over all conditions (Fig. 4E and Material and Methods). The most significant noise propagating factor was H.NS, a general transcriptional repressor that regulates around 5% of all *E.coli* promoters. It belongs to the family of ‘nucleoid associated’ proteins, acting as a histone-like molecule, by binding to curved DNA and inhibiting transcription Dorman, 2004. It is noteworthy that, in eukaryotes, histone positioning plays an important role in determining noise levels Tirosh and Barkai, 2008; Cairns, 2009, although the gene regulatory mechanisms are too different between prokaroytes and eukaryotes to imply a direct mechanistic link between these observations.

The second most significant noise propagating TF is Sigma38 (*rpoS*), which is considered the central regulator of gene expression in stationary phase and under environmental stress, as it interacts with the RNA polymerase and activates genes involved in the overall stress response and genes required to survive long periods of food starvation and growth arrest Tanaka *et al*, 1993; Landini *et al*, 2014. It has been established that, in contrast to rich media, *rpoS* levels in minimal media (the basis of 7 of our 8 conditions) are also high during exponential phase, although the molecular mechanisms behind these differences in *rpoS* activity are unclear Dong and Schellhorn, 2009. Interestingly, it has been reported that in a mutant strain unable to produce ppGpp, noise levels in a set of synthetic genes were significantly reduced in stationary phase compared to a WT strain Guido *et al*, 2007. This observation is in line with the prediction of Sigma38 playing an important role in shaping genome-wide expression noise, as ppGpp is an ‘alarmone’ responsible for regulating genes upon entry into stationary phase Hauryliuk *et al*, 2015 and promoters regulated by Sigma38 require ppGpp for induction Kvint *et al*, 2000. Moreover, it was recently shown that *rpoS* activity is heterogeneous among single cells in M9 glucose Patange *et al*, 2018.

Two further significant noise propagators are CRP and PhoB. CRP is a global regulator of genes involved in carbon source catabolism and its activity has been proposed to reflect carbon source influx You *et al*, 2013. PhoB regulates the response to inorganic phosphate (Pi) starvation and binds to the DNA as a dimer after being phosphorylated by a histidine kinase (PhoR) under Pi limited conditions Santos-Beneit, 2015. We hypothesize that, in our growth conditions, both carbon source influx and Pi concentration are sufficiently limiting that there are significant cell-to-cell fluctuations, leading to fluctuations in the activities of CRP and PhoB. We also note that it has previously been observed that promoters associated with carbon metabolism regulation are overrepresented among high noise promoters Silander *et al*, 2012.

Finally, the two last factors we find to be significantly contributing to noise propagation across all conditions (GadX and GadW), belong to a family of regulators involved in the response to acid stress Tucker *et al*, 2003. The appearance of these factors may also be explained by our experimental setup. Oxygen levels in microtiter plates can easily become limiting and this oxygen deprivation leads to production of fermentation products Salmon *et al*, 2005, even when oxygen is still present Basan *et al*, 2015. Fermentation products are known to acidify the medium Kleman and Strohl, 1994, which can activate the response to acid stress in some cells. Moreover, the fact that we find GadX and GadW as noise propagators is in accordance with a recent publication, where it was shown that heterogeneous expression of the gadBC operon (heavily regulated by GadX and GadW) correlated with single-cell survival to high acid induced by an antibiotic Mitosch *et al*, 2017.

Together our results show that noise propagation by TFs plays a major role in shaping noise levels across genes, and that TFs have different noise propagating activities in each condition, leading to highly condition-dependent noise levels across genes.

### Gene features are organized along two major axes reflecting average expression and regulation

We have shown that, through noise propagation, gene regulation and gene expression noise are intimately coupled, such that highly regulated genes tend to be more noisy and also vary their noise levels more across conditions. We next set out to understand how regulation and expression noise relate to other properties of genes on a genome-wide scale. Previous analysis of gene features has uncovered that, on a genome-wide scale, genes are organized along a one-dimensional axis that relates evolutionary rates, codon bias, and gene expression level Drummond *et al*, 2005, 2006; Drummond and Wilke, 2008; Koonin, 2011, i.e. highly expressed genes tend to have strong codon bias and slowly evolving coding regions, whereas lowly expressed genes tend to have weak codon bias and evolve more rapidly. To investigate how gene regulatory and expression noise properties relate to other gene features we collected a set of features for *E. coli* genes on a genome-wide scale from the literature including the absolute expression levels at both the RNA Taniguchi *et al*, 2010 and protein level Wang *et al*, 2012, sequence properties such as codon bias and evolutionary rates at both synonymous and nonsynonymous sites Drummond and Wilke, 2008 and the number of regulatory inputs of each gene Santos-Zavaleta *et al*, 2019. We then complemented these features with gene regulatory annotations and gene expression features that we measured here including mean expression level, expression plasticity across the 8 growth conditions, mean expression noise, and noise plasticity across the 8 growth conditions.

In total we gathered 10 different gene features and then calculated an overall normalized covariance matrix *C* of correlations between these features, i.e. with *C*_*ij*_ the squared Pearson correlation between features *i* and *j*. We then performed Principal Component Analysis (PCA) of the matrix *C* to characterize the overall genome-wide correlation structure of these gene features. As shown in Suppl Fig. S11, the first two principal components capture significantly more of the total variance than the other 8 components, and more than 50% of the total variance. That is, genes are largely organized along two major PCA axes in the 10-dimensional space of gene features. The first PCA axis sorts genes by their absolute gene expression and evolutionary rate (Fig. 5A). That is, 94% of the weight along this first PCA component is accounted by mean RNA and protein levels, codon bias, and evolutionary rates at synonymous and nonsynonymous sites (Fig. 5A) and, while the absolute expression levels and codon bias are all positively correlated with each other, the evolutionary rates are negatively correlated with these features (Fig. 5C). That is, this first PCA axis recovers the previously observed organization of by their absolute expression levels, codon bias, and evolutionary rates Drummond *et al*, 2005, 2006; Drummond and Wilke, 2008; Koonin, 2011.

**Figure 5:**
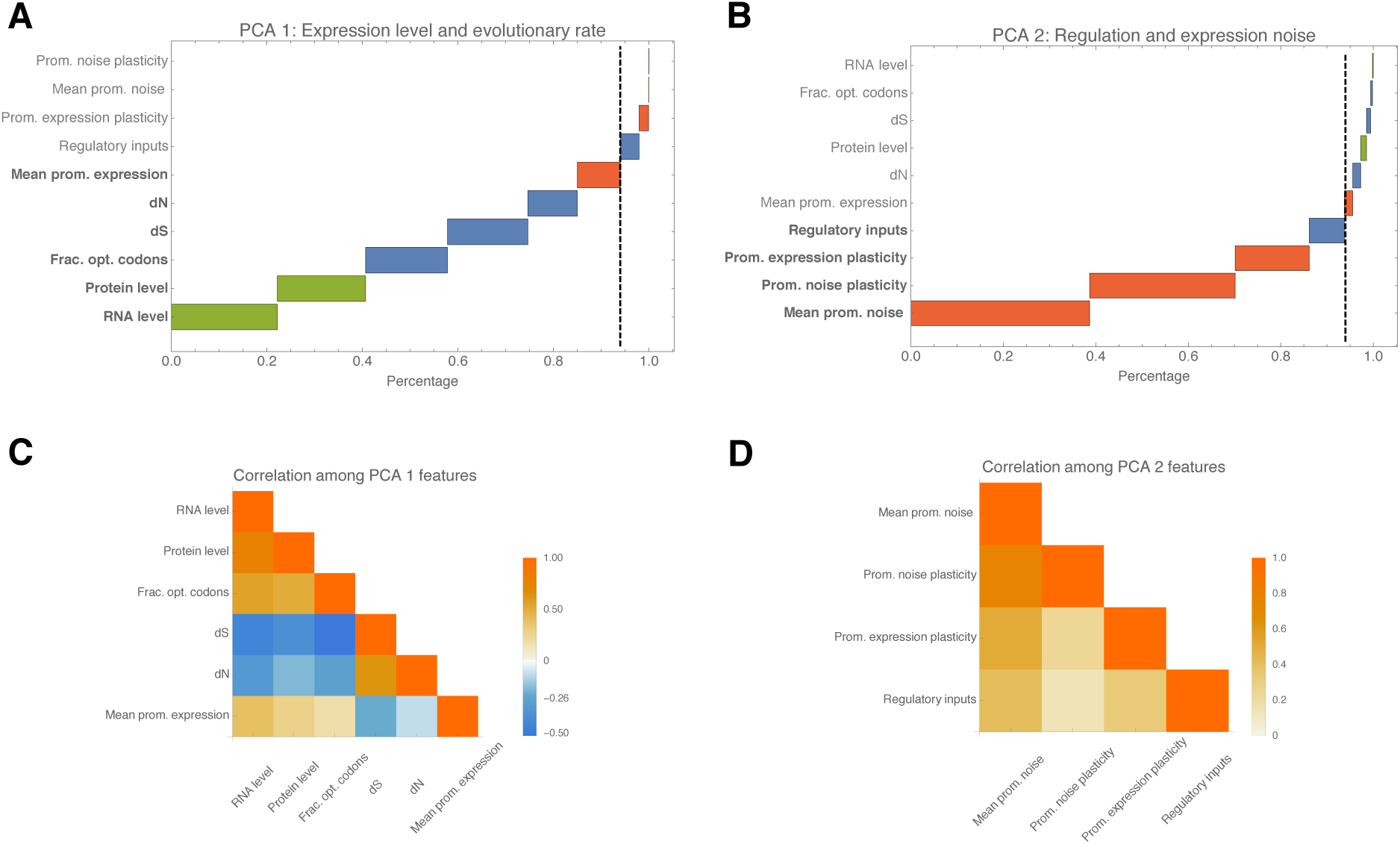
PCA analysis of the genome-wide structure of gene features. **A:** Relative contribution of 10 gene features to the first PCA component, sorted from top to bottom. The features in bold together account for 94% of the vector. In green are expression measurements obtained from previous studies, sequence features are in blue, and features measured in this study are in red. **B:** As in panel A, but now for the second PCA component. **C:** Correlation structure of the features contributing to the first PCA component. Negative correlations are in blue and positive correlations in orange. **D:** As in panel C but now for the second PCA component.

Strikingly, the second PCA axis is almost entirely oriented along features associated with gene regulation and gene expression noise. That is, 94% of the vector’s weight is accounted for by gene expression noise, noise plasticity, plasticity in mean expression, and number of regulatory inputs (Fig. 5B). Moreover, these four features are all positively correlated with each other (Fig. 5D). That is, this second PCA axis organizes genes by their regulation and expression noise. On one end of this axis are constitutively expressed genes that do not change their mean expression level across conditions, and have low noise in all conditions, whereas on the other end of the axis are highly regulated genes that are highly plastic in expression, and have high and varying expression noise across conditions. This result not only further confirms that gene regulation and expression noise are intimately coupled on a genome-wide scale, it also shows that these gene regulatory features are varying *independently* of the absolute expression and evolutionary rate features of the first principal axis.

## Discussion

Although it is now well-established that gene expression is an inherently noisy process, so far little is known in bacteria about how noise levels of genes vary across growth conditions. Here we used high-throughput flow cytometry in combination with a library of fluorescent transcriptional reporters to quantify expression noise of *E. coli* promoters genome-wide. The general picture that emerges from our study is that the expression noise of a given gene in a given condition is the sum of two separate contributions: a minimal amount of noise that derives from global physiological fluctuations and that is approximately equal for all genes, and a highly gene- and condition-specific component that is due to noise propagation from regulators to their targets. Constitutively expressed promoters only exhibit the physiological ‘noise floor’ in each condition, and the more regulated a gene is, the more additional noise from noise propagation it exhibits, and the more variable this additional noise is across conditions.

We observed that the noise floor is itself significantly varying across conditions. In particular, the noise floor systematically decreases with the growth-rate of the cells, and is highest in stationary phase (Fig. 2). Both its dependence on growth-rate, and the fact that this noise floor appears to equally affect all genes, strongly suggest that the noise floor is driven by global physiological fluctuations. However, it is currently unknown what physiological fluctuations most contribute to this noise floor. Fluctuations in chromosome copy number, polymerase concentration, ribosome and charged tRNA concentrations, mRNA decay rates, and fluctuations in growth-rate itself, may all contribute to determining the noise floor. To gain further understanding which fluctuations set the noise floor, and why the noise floor decreases with growth-rate, will likely require quantitative time course data, for example from approaches that combine microfluidics with time-lapse microscopy Wang *et al*, 2010; Kaiser *et al*, 2018.

Our results show that, in addition to this noise floor, each gene exhibits additional expression noise due to noise propagation. This additional noise is not only highly condition-dependent, but different genes show highly diverse behaviors of noise across conditions (Fig. 3). In addition, the more regulatory inputs a gene has, the higher its noise levels tend to be, and the more variable its noise levels across conditions. The intimate relationship between expression noise and gene regulation was further underscored by a global analysis of the correlation structure of a diverse set of gene features. We found that *E. coli* genes are broadly organized along two independent axes in the space of sequence, evolutionary, and gene expression features. While the first axis organizes genes by evolutionary rate and absolute expression level, with low expression and fast evolving genes on one end, and high expression and slow evolving genes on the other, the second axis organizes genes by regulation and expression noise. Here constitutively expressed genes with consistently low expression noise occur on one end of the axis, while highly regulated genes with high expression plasticity, high noise, and high noise plasticity, occur at the other end of the axis.

To identify which TFs are most responsible for noise propagation in each condition we adapted a simple linear model that we previously developed for modeling gene expression in terms of regulatory sites in promoters Balwierz *et al*, 2014, to model gene expression noise in terms of known regulatory inputs and noise propagation activities of TFs. This analysis showed that, in spite of the simplicity of the model, a significant fraction of the variance in expression noise can be explained by noise propagation and we identified both a number of TFs that propagate noise in only a specific condition, and a number of TFs that appear to significantly contribute to noise propagation in all conditions. Among these latter ubiquitously noise propagating TFs are the histone-like TF H.NS, the stationary phase sigma factor Sigma38, the global carbon and phosphate regulators CRP and PhoB, and the Gad TFs involved in acid stress. We strongly suspect that these ubiquitous noise propagating TFs reflect aspects that were shared between all our growth-conditions, i.e. batch culture growth in microtiter plates.

Although our simple linear noise propagation model captures a significant amount of the variation in noise levels, it only captures a modest fraction of the total variance in absolute terms. This is not surprising. In order to make quantitatively accurate predictions of the expression noise of promoters much more realistic models would be needed that take into account that different TFs compete for binding at promoters, that binding rates depend on TF concentrations in a non-linear manner, that interactions between bound TFs and RNA polymerase depend on the relative positioning of sites, and so on. To develop such quantitative models one would likely need more detailed data on the expression dynamics of different promoter architectures. A particularly interesting question that such more detailed data might answer is whether the noise propagation results mainly from fluctuations in TF concentrations across cells, or whether the main source of noise propagation is the stochastic binding and unbinding of the TFs to the promoters.

## Materials and methods

### Strains

All strains used in this study have been previously described Zaslaver *et al*, 2006; Silander *et al*, 2012: each strain carries a transcriptional fusion of a given native *E.coli* promoter followed by a strong ribosomal binding site and *gfp-mut2* (a fast maturing GFP) on a low copy-number plasmid (pUA66 or pUA139 with pSC101 origin, ~ 5 copies per cell). The library contains a construct for ~75% of all intergenic regions longer than 40bp in *E.coli*’s genome flanked by 50 (resp. 150bp) of the downstream (resp. upstream) sequence in order to include most regulatory interactions found on the chromosome.

### Growth conditions

The strains library was stored at −80°C in LB + 7.5% glycerol in microtiter plates. Individual plates were inoculated into fresh media of interest (200 *µ*l) and incubated for two overnights in the same condition before fluorescence measurements. Dilutions (~1/2000) between overnights were done using a 96 Solid Pin Replicator (V&P, 409). The library was grown in a total of 8 different conditions: minimal media, M9 (1mM CaC*l*_2_, 100*µ*M MgS*o*_4_, 1 x M9 salts [Sigma M6030]) supplemented with either 0.2% glucose (w/v), 0.2% glycerol (v/v), 0.2% lactose (w/v), 0.4M NaCl (+ 0.2% glucose [w/v]) or 1.5 ng/ml ciprofloxacin (+ 0.2% glucose [w/v]); a MOPS based synthetic rich media (Teknova, M2105) supplemented with 0.2% glucose, and two stationary phase conditions, where plates were grown for either 16h or 30h in M9 minimal media + 0.2% glucose (w/v). Note that optical density typically saturates after about 10 hours of growth in these conditions (Suppl. Fig. S1).

All media, except the one containing ciprofloxacin, were supplemented with 50 *µ*g/ml kanamycin. The overnights for the sub-MIC ciprofloxacin condition were done in M9 glucose 0.2%, and only at the day of quantification ciprofloxacin was added. On the quantification day, cells were diluted between 200 and 1000-fold depending on the condition (Supplementary Table S1) and grown until mid-exponential phase at 37°, shaken at 600rpm. Growth rates were estimated independently for individual strains in each condition by monitoring the optical density (OD_600_) every 90s during 15-25 hours at 37°C in a plate reader (Biotek Synergy 2). We defined the growth rate as *α* as the slope of a straight-line fit of log(*OD*_600_) against time.

To estimate cell sizes, a strain of the library containing a plasmid without promoter was selected and grown as described. Cells were then placed on a 1% agarose pad and phase contrast images were obtained with a Nikon Ti-E microscope using a 100 × Ph3 objective (NA 1.45) and an Hamamatsu Orca-Flash 4.0 v2 camera. Cell outlines were identified using a custom MATLAB pipeline.

### Flow cytometry quantification of fluorescence

We measured the distribution of GFP fluorescence levels in single cells using a FACSCanto II (BD Biosciences) with a high-throughput sampler (HTS), fluorescence excitation at 488 nm and a 530/30 nm filter for emission. For each strain we collected 5 × 10^4^ events. We used a Bayesian procedure that removes outliers to extract the mean and variance of the log-fluorescence distributions as described in Galbusera *et al*, 2019. Briefly, we first fitted the 4-dimensional signal distribution of forward and side scatter heights and widths by a mixture of a multivariate Gaussian and a uniform ‘background’ distribution. For each event, we then calculated the posterior probability that it derives from the central multi-variate Gaussians, and all events with lower than 50% posterior probability were removed. For the remaining cells, the logarithms of the fluorescence signals (logarithm of the height of the peak) were fitted to a mixture of a Gaussian and a uniform background distribution. That is, the probability of observing log-fluorescence *y* has the form:

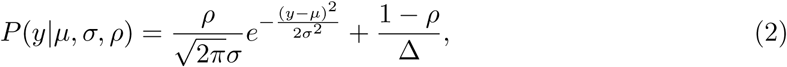

where *µ* and *σ*^2^ are the mean and variance of the log-fluorescence distribution, *ρ* is the fraction of cells deriving from the Gaussian, and ∆ = *y*_max_ − *y*_min_ is the range of observed log-fluorescence values. Given *n* single-cell log-fluorescence measurements *y*_1_, *y*_2_, …, *y*_*n*_ for a given promoter in a given condition, the likelihood is simply given by 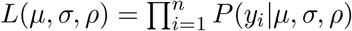 and we fit *µ*, *σ*, and *ρ* by maximizing this likelihood. The data processing method of Galbusera *et al*, 2019 is available as an R package at (https://github.com/vanNimwegenLab/vngFCM.git).

In order to assess reproducibility of the measurements, we measured a subset of the library on multiple days and estimated means and variances of each promoter separately for each day (Suppl. Fig. S4). We defined the mean and variance of each promoter that was measured more than once as the average over its replicates. For each of the individual promoters shown in Fig.3, 6 independent measurements were taken.

### Time-course quantification of fluorescence

One of the plates of the library (95 strains) was grown in M9 + 0.4M NaCl during two overnights. At the day of the quantification a 1/200 dilution was done in 1ml of fresh media in a 96 deep-well plate (with 1 glass-bead per plate for better shaking). The plate was covered with a breathable sealing film and grown at 37°, shaken at 600 rpm. At 9 consecutive time points after dilution (after 0h, 1h, 2h, 3h, 5h, 6.5h, 8.5h, 10h and 11h), 100 *µ*l of the culture was transferred into a 96-well plate and used for fluorescence quantification.

### Minimal variance as a function of mean and noise estimation

We observe a clear lower bound on noise levels (variance of log-fluorescence) that depends on the mean of expression. In previous work Wolf *et al*, 2015 we derived a functional form for this noise floor as a function of mean expression which takes into account that total fluorescence is a sum of background fluorescence and fluorescence deriving from GFP, and that the variance in GFP levels is a sum of a ‘Poissonian’ term that is proportional to mean fluorescence, and a ‘multiplicative’ term proportional to mean fluorescence squared. If we denote the background fluorescence in condition *c* by *f*_*bg,c*_ and the average fluorescence of promoter *p* in condition *c* by 〈*f*_*p,c*_〉, the minimal variance in log-fluorescence takes the form

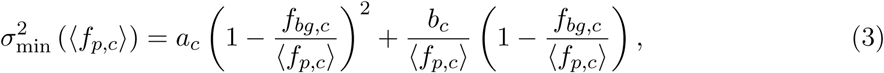

where *b*_*c*_ is the prefactor of the component of the noise proportional to the mean, and *a*_*c*_ is the prefactor of the noise proportional to the square of the mean.

We estimated the average background fluorescence in each condition from plasmids without a promoter upstream of *gfp-mut2* that were included in each individual plate. As the model breaks down in the regime where promoters display fluorescence levels close to background fluorescence, we only considered promoters with mean larger than 2 × 〈*f*_*bg,c*_〉 for further analysis. We fitted the following parameters for the minimal variance in each condition:

| Condition | $a_c$ | $b_c$ | $f_{bg,c}$ | % of promoters above background |
| --- | --- | --- | --- | --- |
| Synthetic Rich | 0.015 | 410 | 180 | 40.0 |
| Ciprofloxacin 1.5ng/ml | 0.05 | 570 | 230 | 58.5 |
| M9 glucose | 0.05 | 530 | 220 | 54.6 |
| M9 lactose | 0.063 | 550 | 220 | 57.2 |
| M9 glycerol | 0.065 | 580 | 205 | 59.0 |
| M9 0.4M NaCl | 0.065 | 500 | 205 | 51.2 |
| Stationary phase 16h | 0.075 | 600 | 190 | 58.2 |
| Stationary phase 30h | 0.075 | 600 | 190 | 58.9 |

To obtain a noise level for each promoter that does not systematically depend on mean, we defined the noise *N*_*pc*_ of promoter *p* in condition *c* as the difference between the measured variance and the fitted minimal variance:

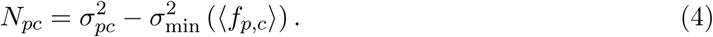

### Noise propagation features

We used the same promoter annotation as in Wolf et al. 2015, where the promoter fragments had been re-annotated by mapping the primer pairs used to construct the library to the *E.coli* K12 MG1655 genome. From all measured promoters we were able to annotate 94% unambiguously to an immediately downstream gene. We obtained all gene-TF regulation annotations from RegulonDB Santos-Zavaleta *et al*, 2019 and counted for each gene the number of unique transcription factors known to regulate it.

We sorted all annotated genes by their average noise across all conditions 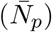 and as a function of a cut-off in 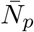, we calculated the mean and standard-error of the number of regulatory inputs of all genes with 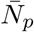 values above the cut-off. We also performed this analysis separately in each of the conditions *c*, using the noise levels *N*_*pc*_ ass opposed to the averages (Suppl. Fig. S9). As a measure of noise plasticity of each promoter *p*, we calculated the variance of the noise levels *N*_*pc*_ across conditions.

### Fitting noise in terms of regulatory inputs

To model noise in terms of regulatory inputs we adapted a method that was previously developed in our group Suzuki *et al*, 2009; Balwierz *et al*, 2014, called Motif Activity Response Analysis, which models gene expression levels in terms of computationally predicted regulatory sites in promoters and condition-dependent activities regulators using a linear model. As explained in the main text, we model the noise *N*_*pc*_ of each promoter *p* in each condition *c* as a linear function of the condition-dependent noise-propagating activities *A*_*rc*_ of the regulators known to regulate promoter *p*, i.e. equation (1).

The binary matrix of regulatory interactions *S*_*pr*_ was constructed using the RegulonDB data Santos-Zavaleta *et al*, 2019, where *S*_*pr*_ = 1 when transcription factor *r* is known to target promoter *p*, and *S*_*pr*_ = 0, otherwise. *N*_*pc*_ is normalized by subtracting the average noise 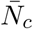 in the condition and *S*_*pr*_ is normalized by subtracting the average of *S*_*pr*_ over all promoters, i.e. the fraction of promoters targeted by regulator *r*.

The noise term, which reflects the deviation between the measurements and our simple model, is assumed to be Gaussian distributed with unknown variance. To avoid overfitting, the model also includes a Gaussian prior over noise-propagation activities *A*_*rc*_ that has mean zero and a variance that is set using cross-validation. In particular, maximal posterior noise-propagation activities *A*_*rc*_ are inferred on 80% of the promoters, and the variance of the prior is set so as to minimize the squared-error of the predictions on the remaining 20% of the promoters. Thus, a different prior is fitted for each condition. As a simple measure of the quality of the fit, we used the fraction of the total variance in the data that is explained by the model (FOV).

For each regulator and condition, we obtain the full posterior distribution over the noise-propagation activity *A*_*rc*_ and use the standard-deviation *δA*_*rc*_ of this posterior as an error-bar for the inferred activity *A*_*rc*_. In addition, we use the *z*-like statistic *z*_*rc*_ = *A*_*rc*_/*δA*_*rc*_ as a measure of significance of regulator *r* in condition *c*.

We defined the average noise-propagating strength *Ā*_*r*_ of each regulator *r* as a weighted average over the 8 conditions:

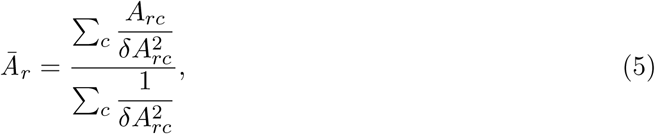

and the corresponding error-bar *δĀ*_*r*_ as

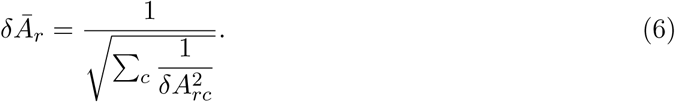

Finally, the average significance 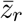 of motifs over all conditions was then estimated as:

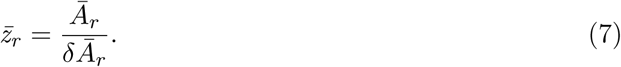

Note that, roughly speaking, *z*_*r*_ corresponds to the number of standard-deviations the activity of regulator *r* is away from zero on average.

To confirm the significance of our fits, we performed tests in which we randomly shuffled the rows of the noise-level matrix *N*_*pc*_, thereby randomizing the association between noise levels and regulatory inputs. We fitted the model to this randomized data and found consistently low FOVs (Figure 4C, yellow bars).

### Principal component analysis

For each promoter we gathered a list of 10 features associated with the immediately downstream gene using both the measurement in this study as well as previously published data. In particular we obtained for each promoter:

1. Average RNA level (data taken from Taniguchi *et al*, 2010).
2. Average protein level (data taken from Wang *et al*, 2012).
3. Fraction of optimal codons (data taken from Drummond and Wilke, 2008).
4. Substitution rate at synonymous sites dS (data taken from Drummond and Wilke, 2008).
5. Substitution rate at non-synonymous sites dN (data taken from Drummond and Wilke, 2008).
6. Average of the mean in log-expression across conditions (this study).
7. Expression plasticity, i.e. variance of the mean in log-expression across conditions (this study).
8. Average of the promoter noise across conditions (this study).
9. Noise plasticity, i.e. variance of the promoter noise across conditions (this study).
10. Number of regulatory inputs (data taken from Santos-Zavaleta *et al*, 2019).

Using these measurements, we calculated a covariance matrix containing all the variances of each of these features across genes, and the covariances of each pair of features. Note that not all features were available for all genes so that, for each pair of features, we estimated the covariance from the set of genes for which both features were available. We then normalized the covariance matrix by dividing each entry *C*_*ij*_ by the square-root of the product of variances, i.e. 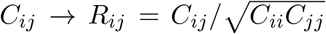, turning it into a matrix of correlation coefficients. We then performed PCA on this correlation matrix. Finally, for the first two principal components we calculated what fraction of the principal vector’s length was accounted for by each feature.

## Acknowledgements

We would like to thank our FACS Core Facility, specially Janine Bögli, for assistance with flow cytometry, Guillaume Witz for support with microscopy analysis and Mikhail Pachkov for help with setting up MARA. Thanks to Urs Jenal, Gasper Tkačik and Olin Silander for useful discussions on the project; and to Dany Chauvin and Ðorđe Relic for comments on the manuscript. This work was partly funded by the Werner Siemens Stiftung through a fellowhip to AU, the SystemsX.ch StoNets grant, and the SNF grant 31003A_159673 to EvN.

## Author contributions

EvN and TJ designed the study. AU performed all experiments, except fluorescence quantification with flow cytometry in the Ciprofloxacin condition, which was done by GB. LG wrote the R package for the estimation of mean and variances from flow cytometry data. AU, TJ and EvN analyzed the data. AU, TJ, and EvN wrote the paper.

## Supplementary information

**Table S1:** List of conditions: Description of the 8 environmental conditions in which the library of native *E.coli* promoters Zaslaver *et al*, 2006 has been grown. The chosen environmental conditions comprised a MOPS based synthetic rich media, minimal media (M9) with different carbon sources (glucose, lactose and glycerol), an osmotic and DNA damage stress (0.4M NaCl and Ciprofloxacin 1.5 ng/ml both supplemented with glucose), as well as two time points in stationary phase (16 and 30 hours of growth in M9 glucose).

| Condition | Condition during overnights | Hours of growth before FACS measurements | Dilution after ON | Growth rate +/- sd (h<sup>-1</sup>) | Mean area +/- sd (in $\mu\text{m}^2$) |
| --- | --- | --- | --- | --- | --- |
| Synthetic Rich* | same | 2h | 1/1000 | 1.59 +/- 0.28 | 4.3 +/- 1.4 |
| M9** + Ciprofloxacin 1.5 ng/ml (+ 0.2% glucose) | M9 + 0.2% glucose | 4h | 1/200 | 0.69 +/- 0.07 | 2.7 +/- 1 |
| M9** + 0.2% glucose | same | 4h | 1/200 | 0.67 +/- 0.05 | 2 +/- 0.6 |
| M9** + 0.2% lactose | same | 4h | 1/200 | 0.58 +/- 0.03 | 2.1 +/- 0.6 |
| M9** + 0.2% glycerol | same | 5h | 1/200 | 0.5 +/- 0.11 | 1.6 +/- 0.5 |
| M9** + 0.4M NaCl (+ 0.2% glucose) | same | 15h | 1/500 | 0.37 +/- 0.03 | 1.4 +/- 0.3 |
| Stationary phase 16h (M9** 0.2% glucose) | same | 16h | 1/200 | x | 1.3 +/- 0.4 |
| Stationary phase 30h (M9** 0.2% glucose) | same | 30h | 1/200 | x | x |

\* MOPS based. Commercially available (Teknova M2105)
\*\* Prepared as follows: 1mM CaCl<sub>2</sub>, 100 $\mu$ M MgSO<sub>4</sub>, 1 x M9 salts (Sigma M6030), 50 $\mu$ g/ml Kanamycin, dH<sub>2</sub>O

**Figure S1:**
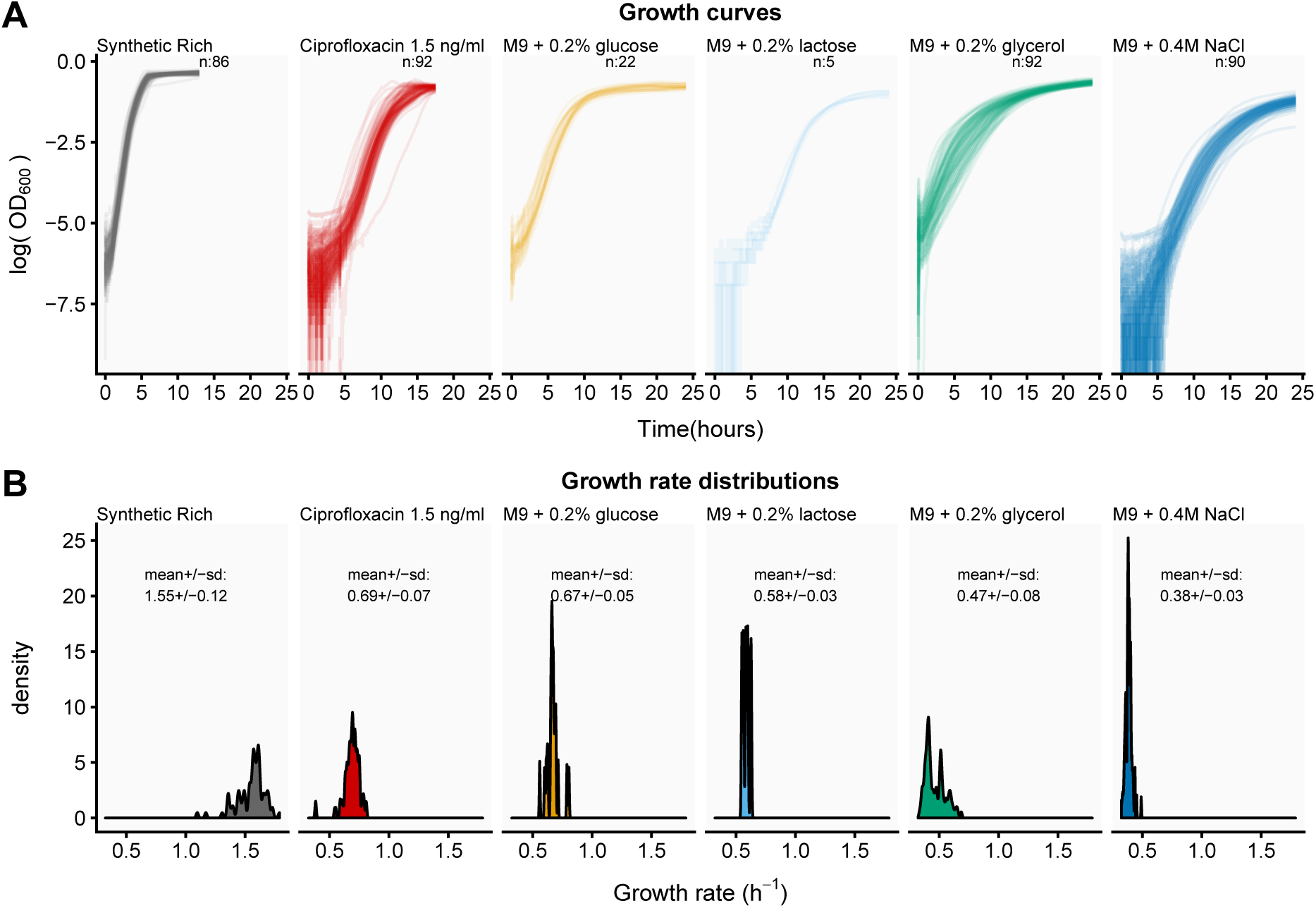
Growth curves across conditions. **A:** *OD*_600_ (log-scale, y-axis) as a function of time (in hours, x-axis). We measured *OD*_600_ in individual strains growing in bulk at intervals of 90 seconds during 15 to 25 hours. The number of strains used per condition is indicated in each panel. **B:** Density distribution of the estimated growth rates in each condition. The growth-rate *α* was defined as the slope of a linear fit of log(*OD*_600_) against time.

**Figure S2:**
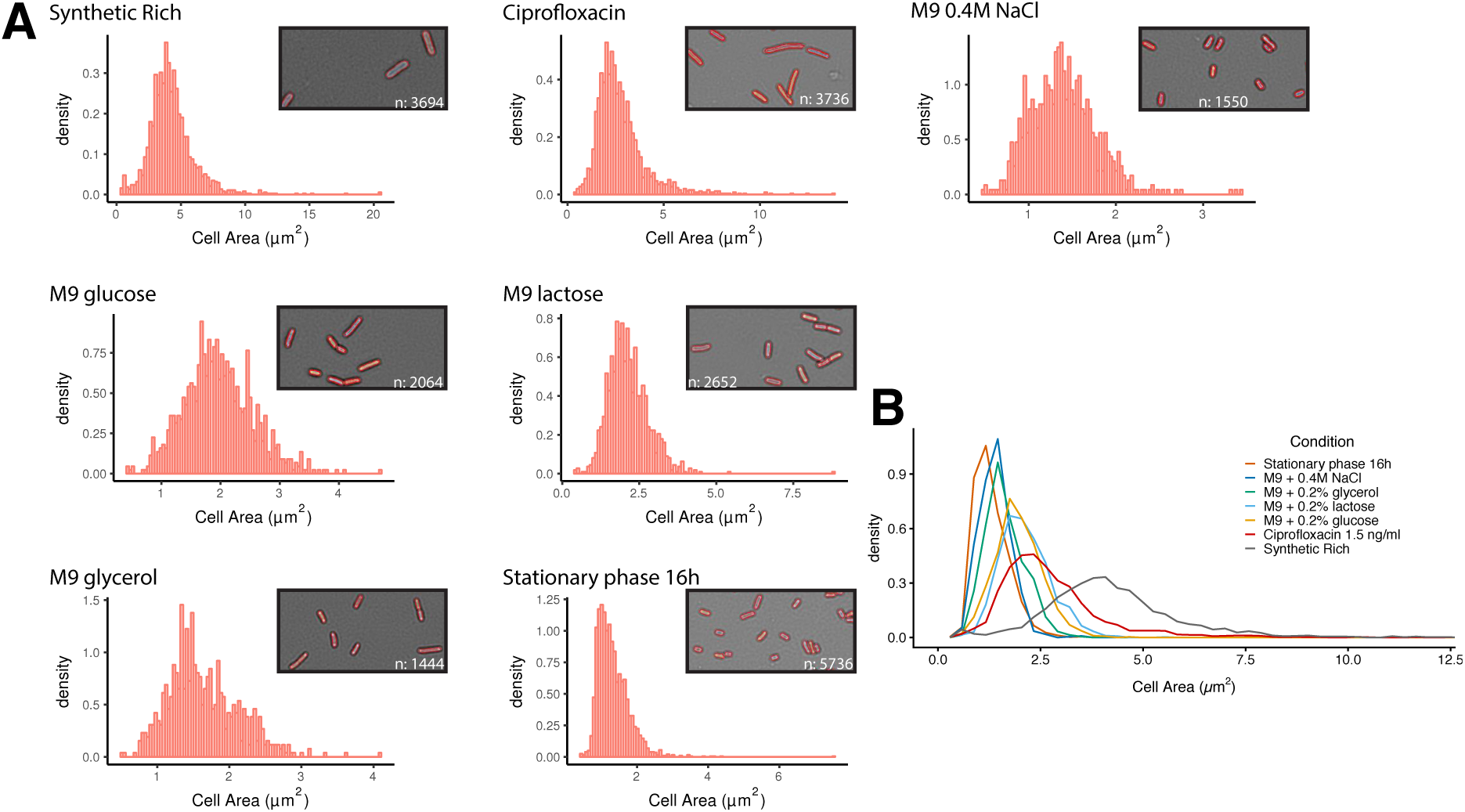
Cell sizes distributions. **A:** Histograms of the distribution of single-cell areas (*µm*^2^, x-axis) in each condition. The insets in each condition show segmentation examples together with the number of cells used to estimate the mean and standard-deviation of the areas. **B:** Kernel-density estimates of the distribution of areas across all conditions (Areas bigger than 12.5 *µm*^2^ are not shown).

**Figure S3:**
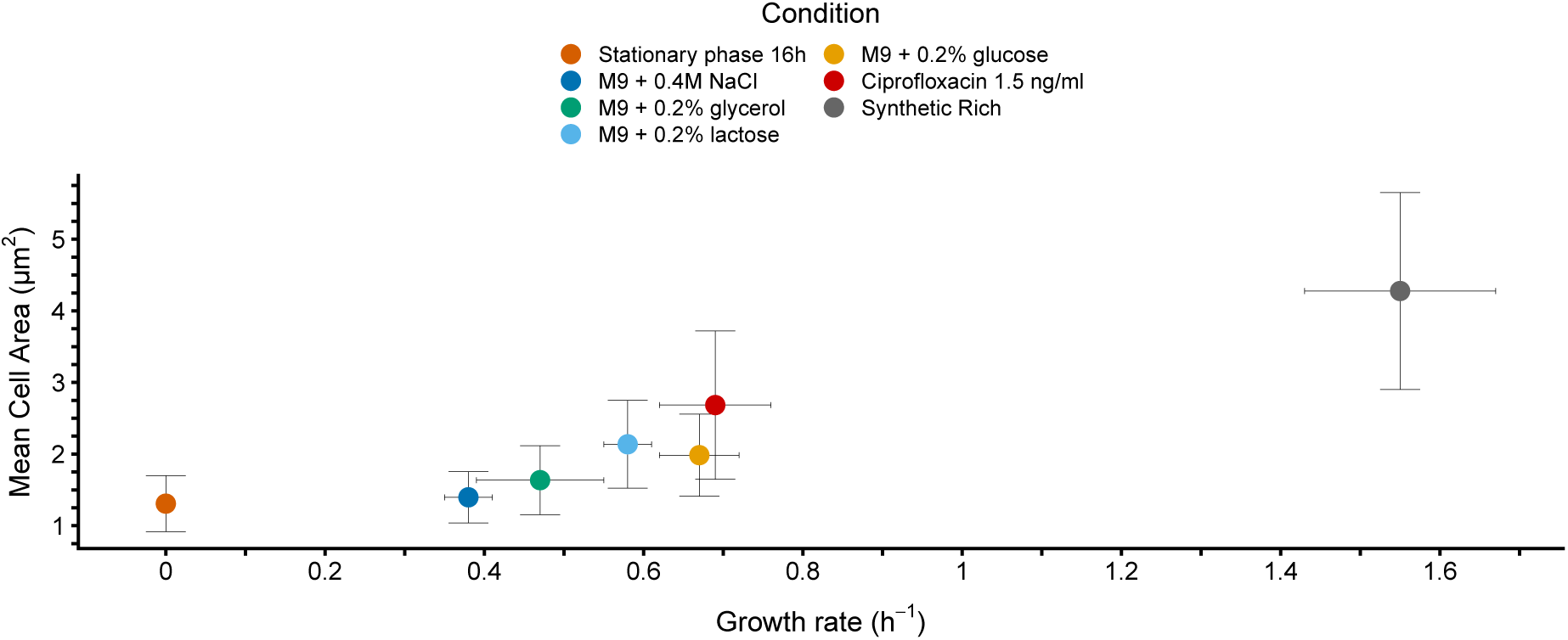
Cell area as a function of growth rate. Mean cell area (in *µm*^2^) as a function of the growth rate (*h*^−1^) in all conditions except stationary phase 30h. Cross-hairs indicate standard-deviations.

**Figure S4:**
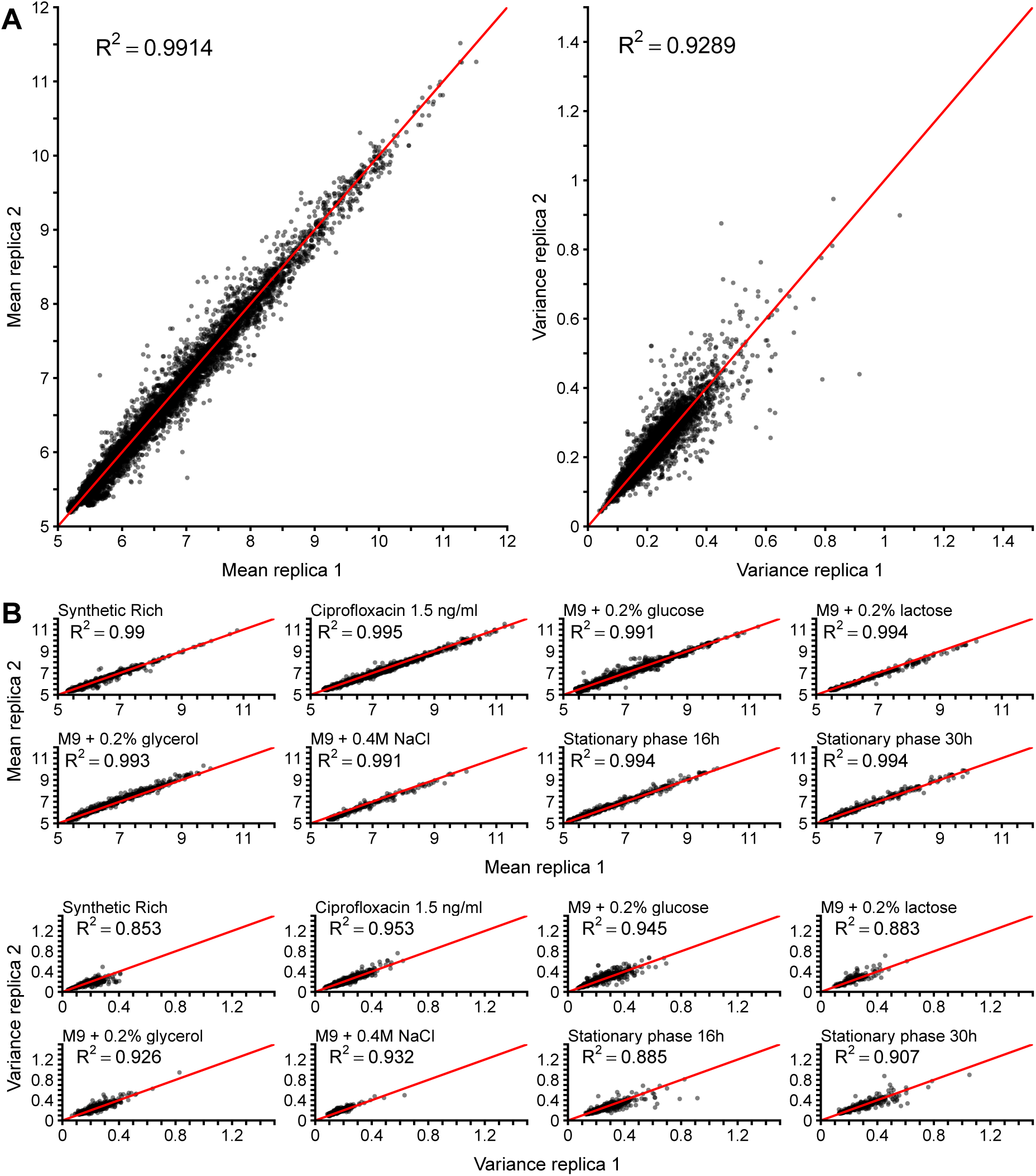
Reproducibility of measured means and variances. **A:** Means (left-panel) and variances (right-panel) of promoters (each represented by a black dot) measured on different days. The Pearson squared-correlations are indicated in each panel. **B:** Reproducibility of means (top panel) and variances (bottom panel) separately for each condition. Pearson squared correlations are indicated in each panel.

**Figure S5:**
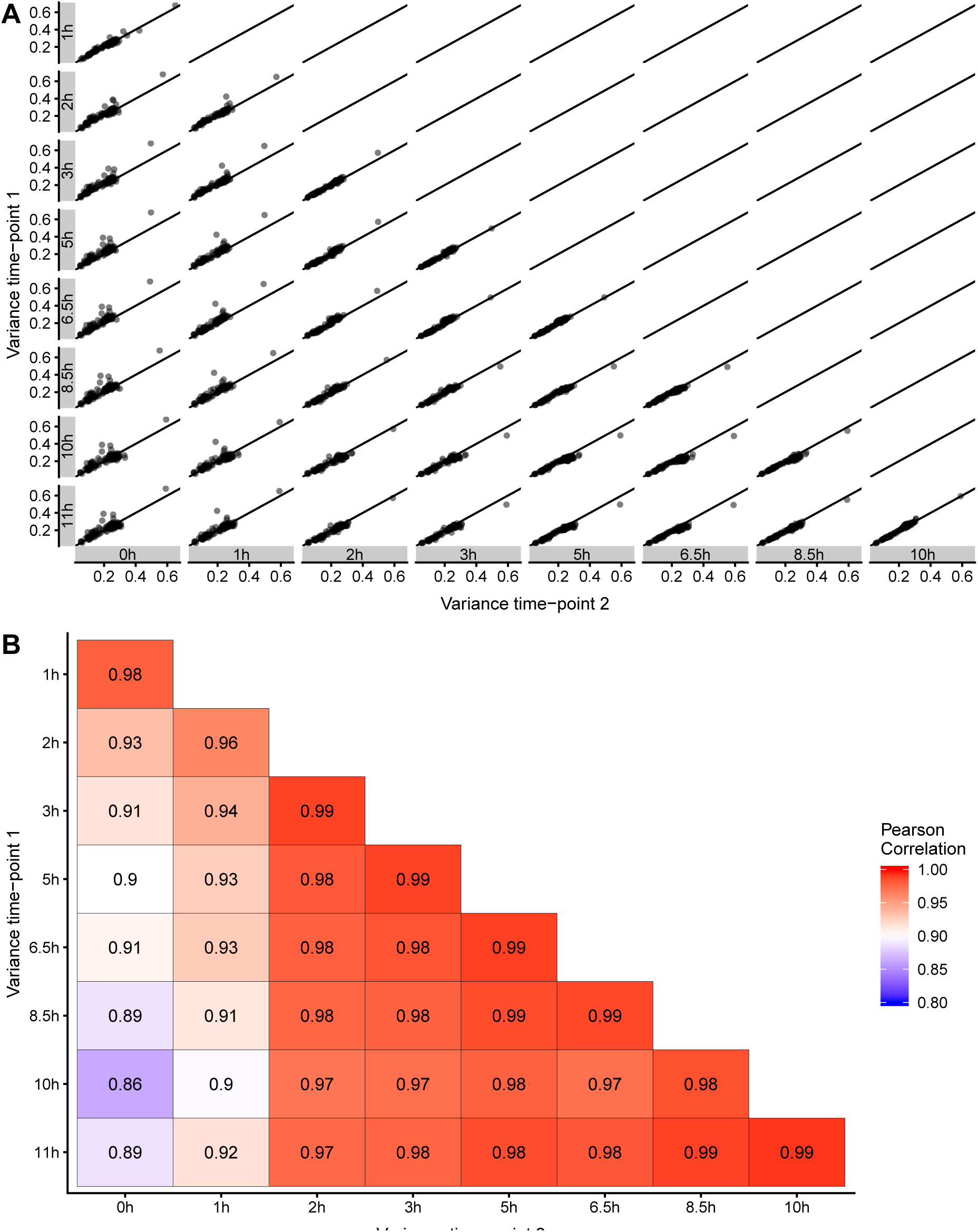
Reproducibility of measured mean fluorescences at different time-points during growth. **A:** Correlations of mean expression levels for 95 promoters from the library, measured at consecutive time points during growth in M9 + 0.4M NaCl (+ 0.2% glucose). The time points range between 0h (freshly diluted culture) and 11 hours. The grey boxes on the axes indicate the time points that are being compared. **B:** *R*^2^ Pearson correlation coefficients of measured mean expression levels for all pairs of timepoints.

**Figure S6:**
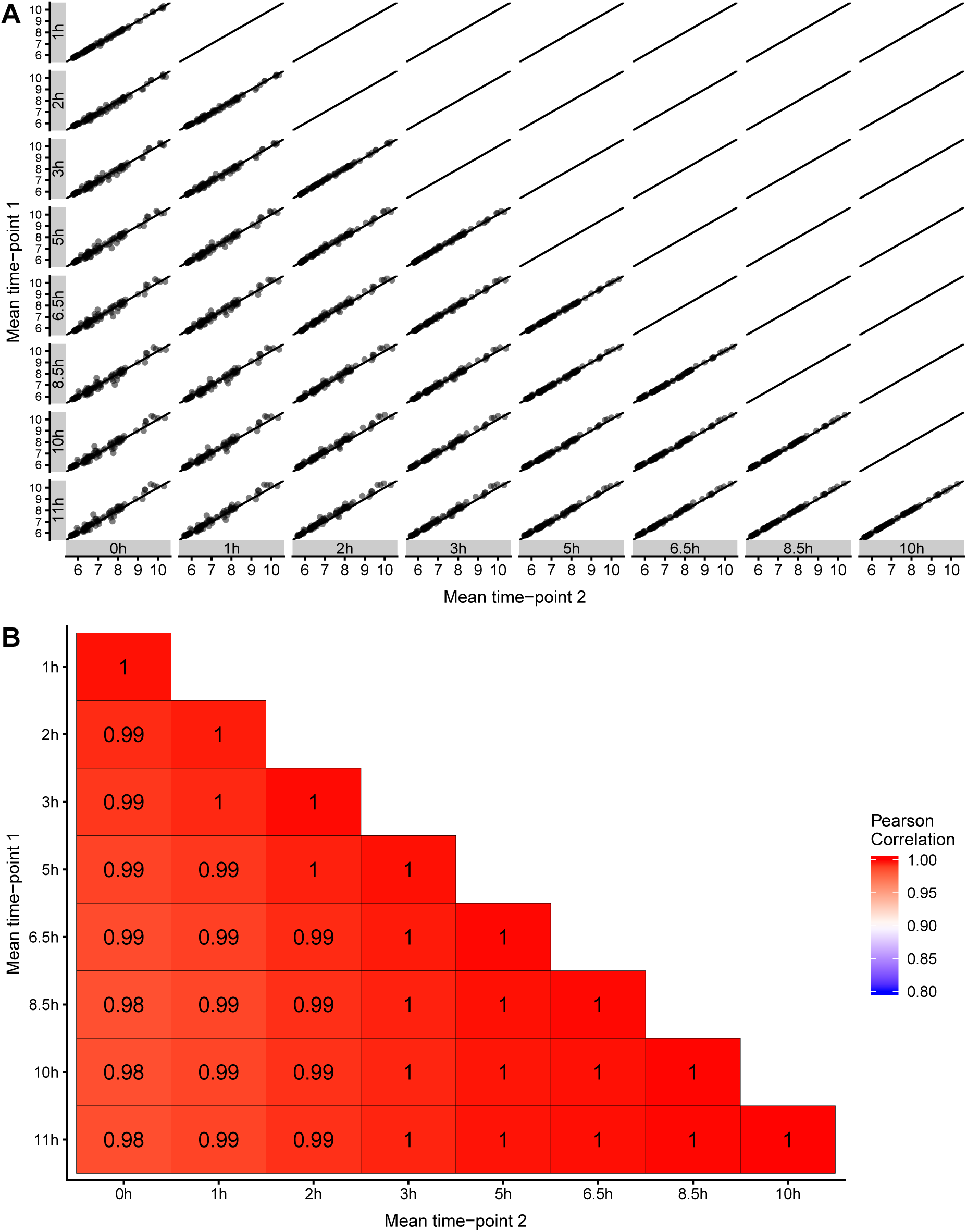
Reproducibility of measured fluorescence variances at different time-points during growth. **A:** Correlations of variance in expression levels for 95 promoters from the library, measured at consecutive time points during growth in M9 + 0.4M NaCl (+ 0.2% glucose). The time points range between 0h (freshly diluted culture) and 11 hours. The grey boxes on the axes indicate the time points that are being compared. **B:** *R*^2^ Pearson correlation coefficients of measured variances in expression levels for all pairs of timepoints.

**Figure S7:**
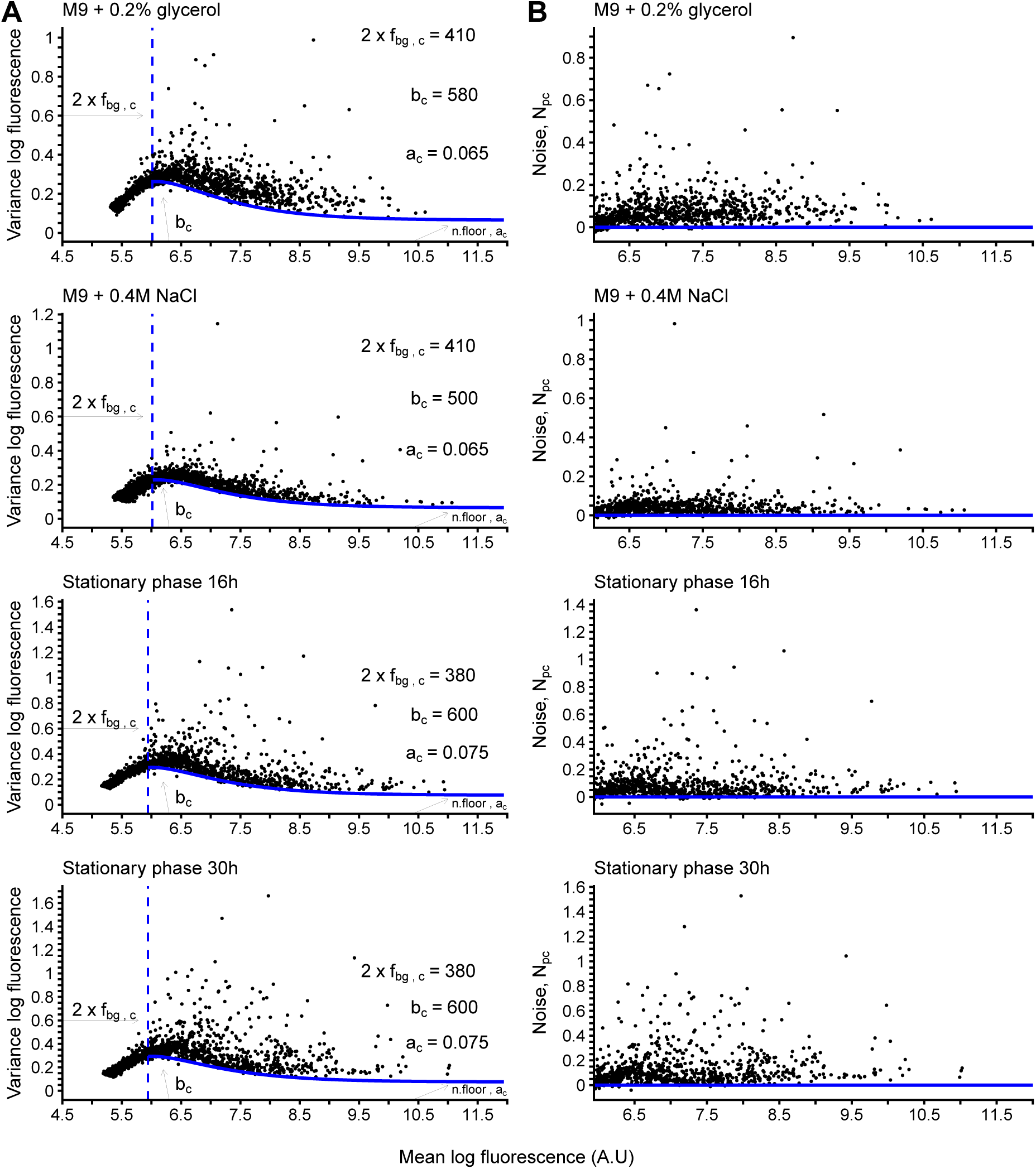
Means and variances of promoters from the library in all conditions. **A:** Variance as a function of mean for all promoters measured in each condition. Each promoter is represented by a black dot. The blue line indicates the predicted minimal variance as a function of mean. The model breaks at fluorescence levels close to background (left of the vertical blue dashed line), thus we only considered promoters above it. The number of promoters measured per condition is annotated inside each panel. **B:** Noise-level *N*_*pc*_ as a function of mean after correcting for the mean-dependent noise floor, i.e. differences between measured variance and minimal variance (Figure continued on next page).

**Figure S8:**
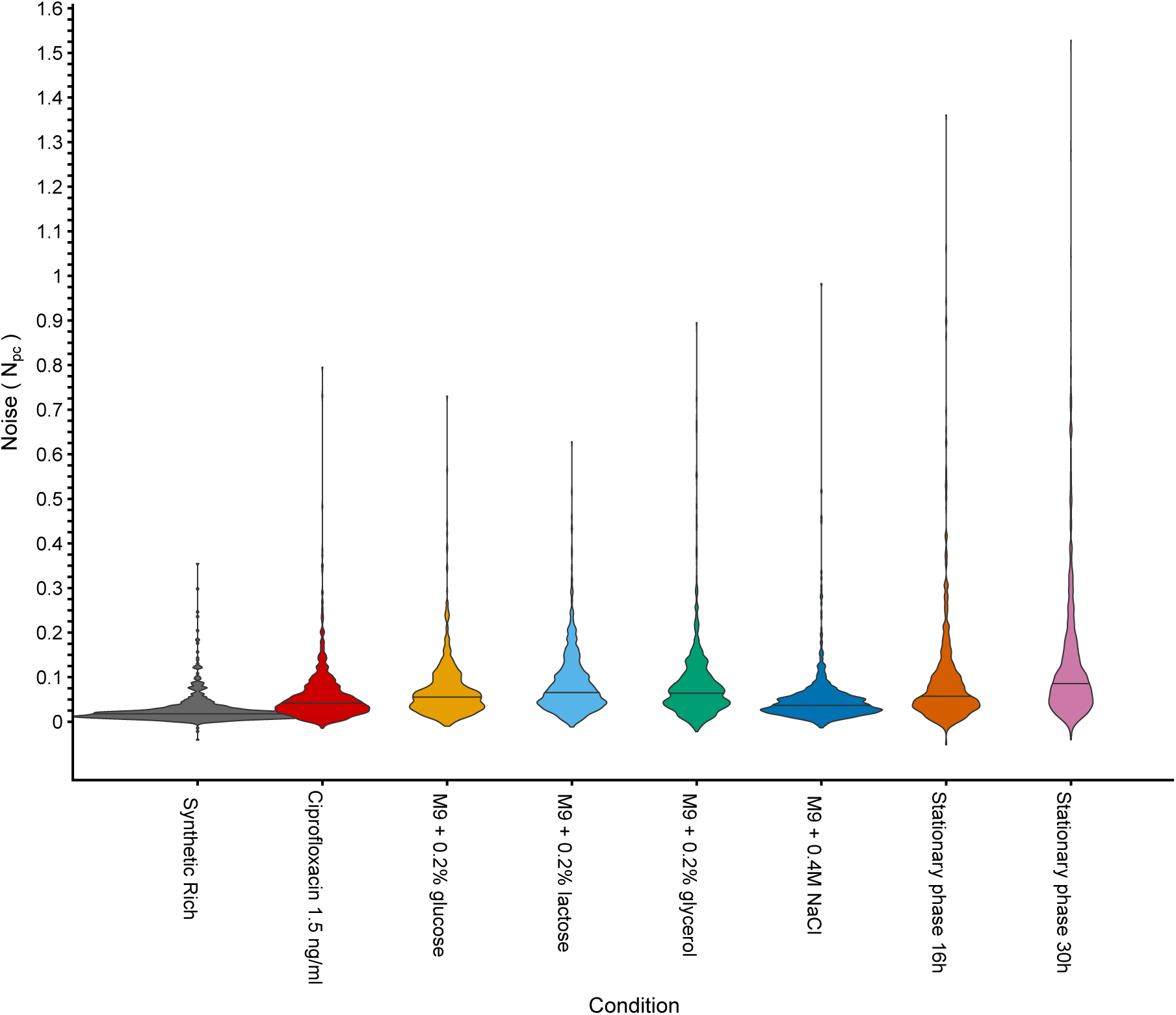
Noise levels across all conditions. The violin plots show the Noise distributions of all promoters in each of the measured conditions. The horizontal lines indicate the medians.

**Figure S9:**
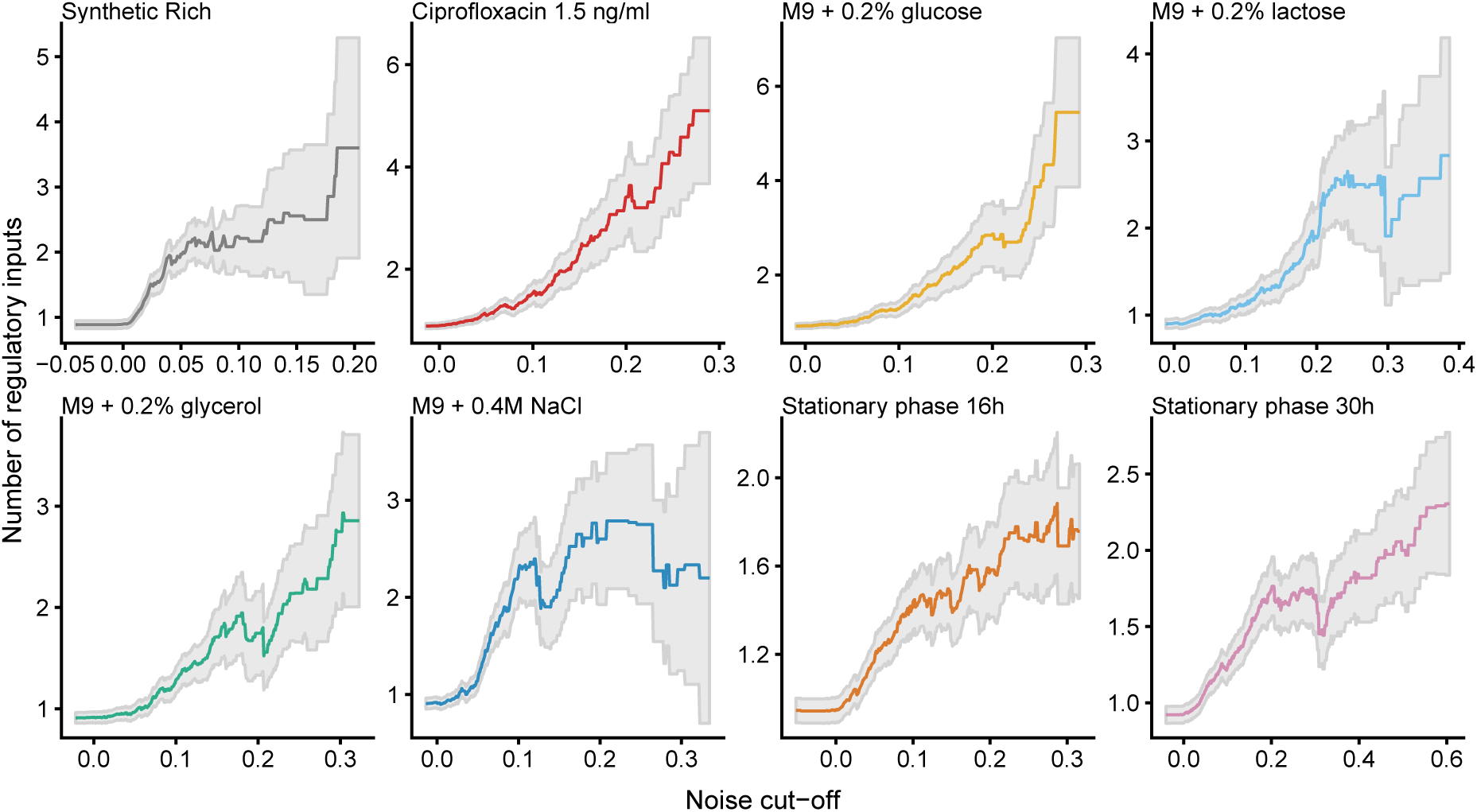
Noise as a function of number of regulatory inputs. In each condition we sorted promoters by their noise level *N*_*pc*_, and calculated the mean and standard-error of the number of known regulatory inputs (y-axis) of all promoters above a cut-off in *N*_*pc*_ (x-axis). Regulatory interactions were annotated from Regulon DB Santos-Zavaleta *et al*, 2019.

**Figure S10:**
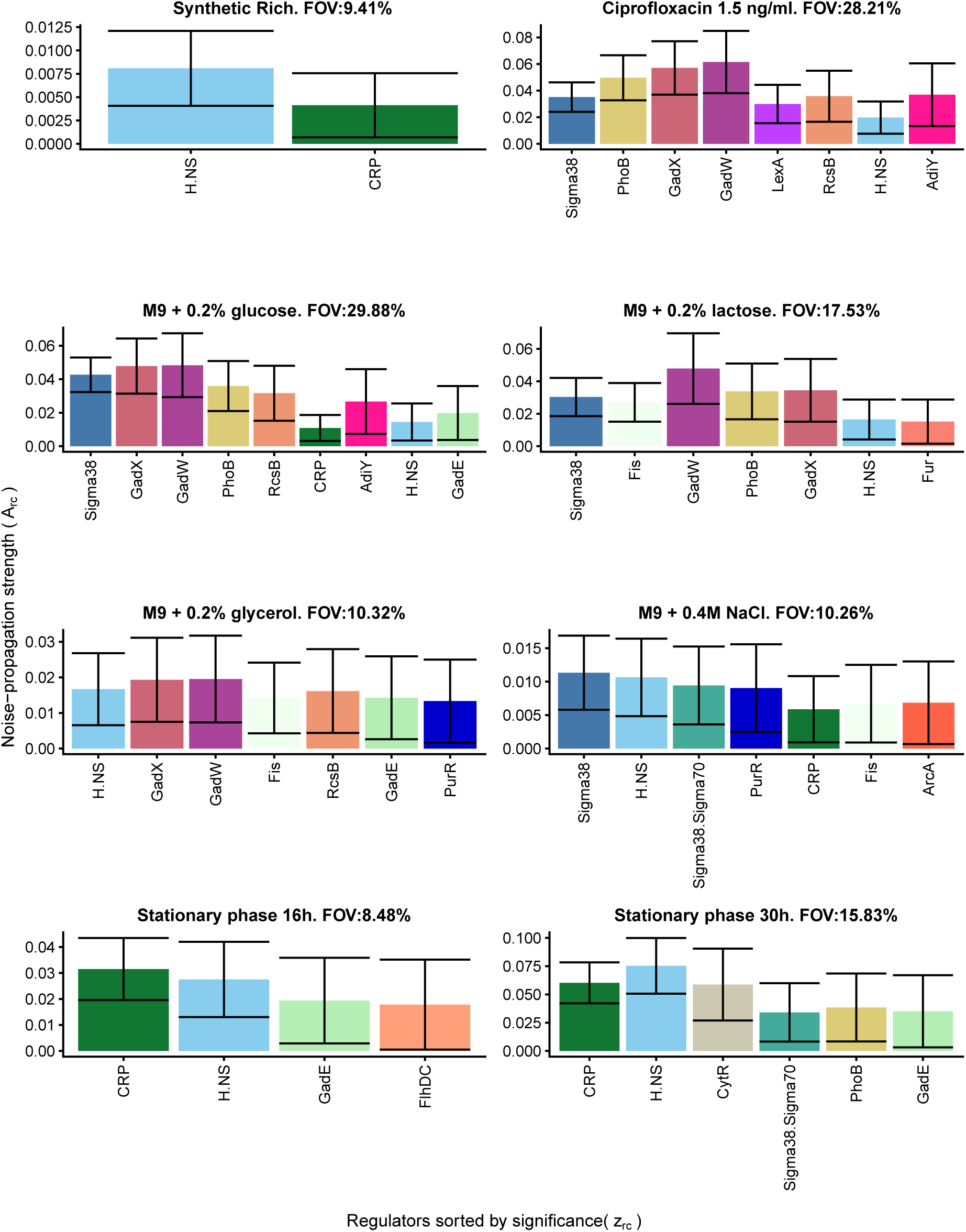
Strongest noise-propagators in each condition. Each panel corresponds to one growth condition and shows the inferred noise propagation strengths *A*_*rc*_ for the transcription factors for which *A*_*rc*_ > *δA*_*rc*_ in that condition. The TFs are sorted by their overall signifiance *z*_*r*_. The condition is indicated above each panel together with the fraction of variance (FOV) explained by the model.

**Figure S11:**
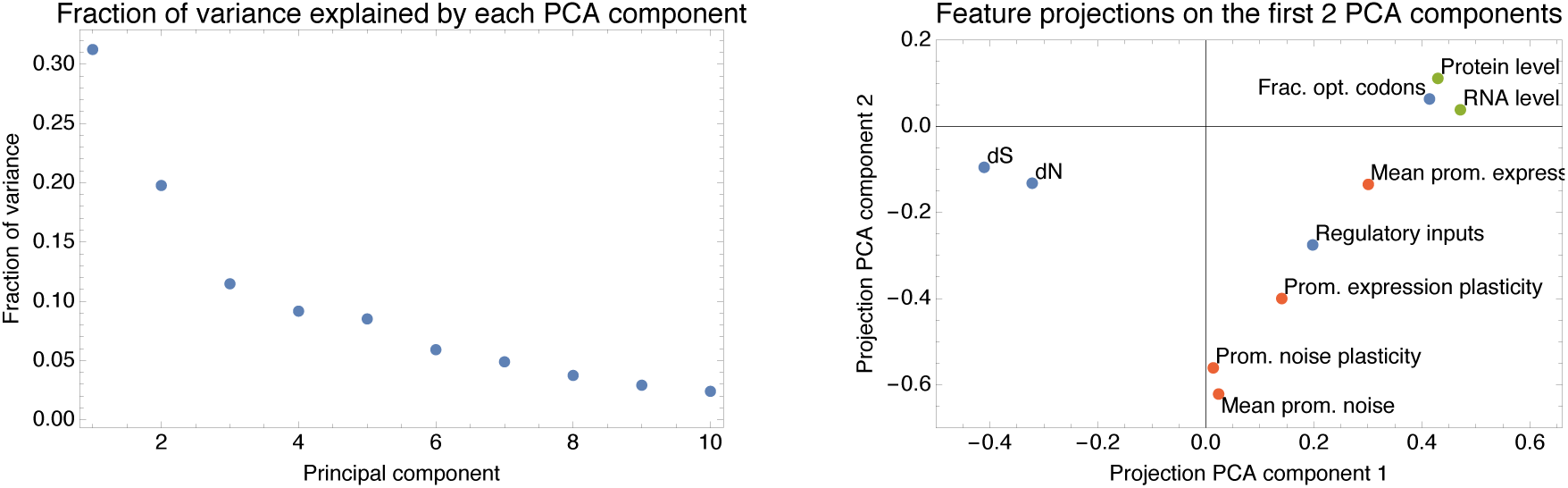
Principal component analysis of the 10 gene features. **A:** Fractions of the total variance in gene features captured by each of the PCA components. Note that the first two components together capture more than 50% of the variance. **B:** Projection of each of the 10 features on the first two PCA components. Expression levels from the literature are shown in green, sequence features are shown in blue, and gene expression features measured in this study are shown in red.

